# Phosphatidylserine controls synaptic targeting and membrane stability of ASIC1a

**DOI:** 10.1101/2022.09.29.509830

**Authors:** Di-Shi Liu, Xing-Lei Song, Ming-Gang Liu, Jianfei Lu, Yu Huang, Jaepyo Jeon, Guofen Ma, Yong Li, Lucas Pozzo-Miller, Michael X. Zhu, Tian-Le Xu

## Abstract

Phospholipid-protein interaction is highly specialized at the membranous nanodomains and critical for membrane receptor signaling. Calcium-permeable acid-sensing ion channel isoform 1a (ASIC1a) is a major neuronal proton sensor that contributes to synaptic plasticity. The functional outcome of ASIC1a is dependent on its surface targeting in synaptic subdomains; however, the lipid environment for ASIC1a and its role in channel targeting remain poorly understood. Here, we report that anionic phosphatidylserine (PS) is enriched in dendritic spines during neurodevelopment and it directly binds to ASIC1a through an electrostatic interaction with a di-arginine motif at ASIC1a C-terminus. PS regulates the membrane targeting and function of ASIC1a, which are both strongly suppressed by inhibition of PS synthesis. In cortical neuron dendrites, both PS and ASIC1a are predominately localized to peri-synaptic sites of spine heads, surrounding instead of overlapping with postsynaptic markers, PSD-95 and GluN1. Uncoupling the interaction between PS and ASIC1a by changing the charges to neutral or acidic at the di-arginine PS-binding motif, or applying a membrane penetrating competing peptide, caused mistargeting of ASIC1a at the synaptic sites, an overall increase in internalization and/or cytoplasmic accumulation of ASIC1a, and a decrease in its channel function. Together, our results provide novel insights on lipid microenvironment that governs ASIC1a expression and function at the membrane surface, especially peri-synaptic regions of dendritic spines, through an electrostatic interaction with anionic phospholipids.

## Introduction

Phospholipids are the major components of bilayer membrane systems in cellular organisms (*1*). Different types of phospholipids have various functions in recruiting membranous proteins, such as enzymes and receptors, and regulating their subcellular distributions and activities (*2–4*). Phosphatidylserine (PS) is a relatively minor constituent of eukaryotic membrane phospholipids and often considered as a classic signal that triggers apoptotic recognition and blood coagulation when exposed on the extracellular surface (*5, 6*). However, under normal conditions, PS preferentially exists in the inner leaflet of the plasma membrane (PM) (*7*), where it makes a comparatively large contribution to the negative surface charges (*8*). PS binds several key membrane-associated proteins such as Ras and Akt, by which it regulates surface anchoring and thereby downstream signaling of these molecules (*9*).

Interestingly, PS is particularly enriched in mammalian brains, where it serves important roles in brain development and function. Both gain and loss of PS Synthase I (PSSI) activity cause intellectual disability, such as Lenz-Majewski syndrome and Nablus mask-like facial syndrome, respectively (*10, 11*). As the fundamental unit of neurotransmission and plasticity that underlies all forms of learning and memory, neuronal synapses are specialized structures with complex lipid and receptor compositions. Recently, it was shown that PS externalization marks the specific synapses for elimination by microglia, a process important for synaptic pruning during development and synaptic loss in neurodegeneration (*12–14*). However, it remains to be elucidated how PS is distributed in the synapses under normal physiological conditions and what roles it plays in addition to serving as a “eat me” signal of the excess or damaged synapses.

Acid-sensing ion channels (ASICs) belong to the amiloride-sensitive degenerin /epithelial Na^+^ channel (ENaC) superfamily, and they are widely expressed throughout the mammalian central and peripheral nervous systems (*15*). The ASIC1a isoform is a key member of the ASIC family in the brain that senses extracellular pH fluctuations under physiological conditions (≤ 7.2) (*16*) to mediate sodium and calcium influx by forming homomeric and heteromeric channels (*17*). Numerous studies have shown that the ASIC1a-containing channels are enriched in neuronal dendritic spines (*18*) and they contribute to spine morphogenesis (*19, 20*), synaptic transmission (*21, 22*), and long-term potentiation (LTP) (*23–25*), indicating that ASIC1a plays important roles in both structural and functional plasticity in the nervous system. At the cellular level, the functionality of ASIC1a largely depends on their PM targeting and proper surface presentation (*18*). Although activity-dependent regulatory pathways for ASIC1a surface delivery have been described,(*26–28*), how synaptic trafficking and localization of ASIC1a are regulated by the lipid membrane microenvironment remains unknown.

In the present study, we show that anionic phospholipids, especially PS, directly bind and stabilize ASIC1a at the synaptic membrane. An electrostatic interaction between the negatively charged phospholipid head group of PS and a positively charged binding motif of ASIC1a is responsible for this lipid-protein recognition. Uncoupling this interaction leads to mistargeting and intracellular retention of ASIC1a, which in turn suppresses its channel function. Our findings support the idea that PS anchors ASIC1a on the peri-synaptic surface of dendritic spines, where the channel senses spillover protons from the synaptic cleft under conditions of high frequency stimulation to facilitate LTP induction.

## Materials and methods

### Animals

All animal procedures were carried out in accordance with the guidelines for the Care and Use of Laboratory Animals of Shanghai Jiao Tong University School of Medicine (Policy Number DLAS-MP-ANIM.01–05). ASIC1a knockout mice (C57BL/6J background) were the generous gifts of Professor Michael J. Welsh (Howard Hughes Medical Institute, University of Iowa). All animals were housed in groups of three to four per cage under standard husbandry conditions (12 hr light/dark cycle, temperature 22–26 °C, humidity 55–60%) with *ad libitum* access to water and food.

### Reagents

Drugs and chemicals were purchased from Sigma-Aldrich unless where indicated otherwise. All peptides were synthesized in GL Biochem Corporation Ltd (Shanghai, China). FITC conjugate of Cholera Toxin B subunit was purchased from Sigma. The ASIC1a inhibitor, PcTX1, was purchased from Peptide Institute (Osaka, Japan). The following antibodies (p for polyclonal and m for monoclonal) were used for immunoblotting: anti-ASIC1a pAb (1:500, Santa Cruz), anti-cMyc mAb (1:1000, AbRC of SIBS, CAS, China), anti-GAPDH mAb (1:3000, KangCheng, China), anti-GFP pAb (1:500, Santa Cruz or Invitrogen), anti-GFP mAb (1:1000, Roche), anti-GST mAb (1:2000, Sigma), anti-α-tubulin mAb (1:3000, Sigma).

### Plasmid Construction

The open reading frames of human ASIC1a and ASIC2a from GW1-CMV-ASIC1a and GW4-CMV-ASIC2a constructs, respectively, were subcloned into the pEGFP-C3 vectors (between XhoI and HindIII digestion sites). pEGFP-C1-Lact-C2 construct was from Sergio Grinstein’s Lab (Addgene). The EGFP protein appended on the N-terminus of ASICs or Lact-C2 (between AgeI and XhoI sites) was also substituted by mCherry (from pAW-mCherry vector) or 6×myc-tag (from pCS2-6mt vector) for specific use. For *in vivo* transfection, fluorescent-protein-tagged ASIC1a and Lact-C2, EGFP and mCherry, were individually subcloned into pAAV-MCS vectors and driven by the human Synapsin I promoter for neuronal expression. pCI-SEP-GluN1 construct was from Robert Malinow’s Lab (Addgene). Plasmids pcDNA3-EGFP-Cdc42 and pcDNA3-EGFP-Rac1 were from Gary Bokoch’s Lab (Addgene). pEGFP-N1-Lamp1 construct was a kind gift of Dr. Yi-Zheng Wang (Beijing Institute of Basic Medical Sciences). Plasmid pCMV-SPORT6-Rab11a (mouse) was purchased from GeneCopoeia Inc. Rab11a was subcloned into the pEGFP-C3 vector between XhoI and HindIII sites, and fused with EGFP, mCherry or 6×myc-tag, respectively. All constructs were confirmed by DNA sequencing, and verified by transfection and subsequent imaging or functional assays.

### Generation of shRNAs

The cDNAs of mouse PS Synthase I (PtdSSI, PSSI) and II (PtdSSII, PSSII) were obtained from GeneCopoeia, Inc. and Thermo Fisher Scientific Inc., respectively. The open reading frames of PSSI and PSSII were subcloned into pEGFP-C3 between XhoI and HindIII sites, and the upstream EGFP was then substituted with 6×myc-tag. After myc-mPSSI and myc-mPSSII were transfected into CHO cells, their normal expression and molecular weights were further verified by immunoblotting. Four shRNA candidates targeting different sequences for each of the two genes and a negative control (NC) oligo were designed, synthesized, and inserted into pLKD-RFP-U6-shRNA vector (Neuron Biotech Co., Ltd). The nucleotide target sequences are as follows: shPSSI-#456, 5’-GCC TTG TTG ATC CGT AGT TAT; shPSSI-#457, 5’-GCA ACC ACG AAA GCC ATT CTT; shPSSI-#458, 5’-GAA TCG TTA GTC ACT TTG ATA; shPSSI-#459, 5’-GCA GTT GAC TGA GTT GAA TAC; shPSSII-#598, 5’-GCC GAC AGT TTC TGA AGT ATG; shPSSII-#599, 5’-GCA CAC TTC ATT GGC TGG TAT; shPSSII-#600, 5’-GCT GTC CCT GAA GAC ATA TAA; shPSSII-#601, 5’-GGA ACA TTC CAA CCT ACA AGG; shRNA-NC, 5’-TTC TCC GAA CGT GTC ACG T. All constructs were sequenced, and knockdown efficiencies were further determined by co-transfection with myc-tagged PSSI/II proteins and immunoblotting with anti-myc antibody. Neuron Nucleofector kit (Amaxa) was used for shRNA electroporation when necessary, according to the manufacturer’s instructions.

### Site-directed mutagenesis

In order to change, insert or delete amino acids in different constructs, site-directed mutagenesis or In-fusion (Clontech) procedure was performed. R467 and R468 of ASIC1a were mutated to alanine, glutamine, glutamate, or lysine, respectively, to change the electrostatic properties of the di-basic PS binding motif. The HA-tags (hemagglutinin epitope, YPYDVPDYA) were inserted into the different sites of the coding sequence of ASIC1a ectodomain and detected with anti-HA antibody to screen for the ASIC1a construct with the highest efficiency of surface labeling (*18*). Residues W26, W33 and F34 of Lact-C2 were mutated to alanine to obtain the PS-non-binding mutant (Lact-C2-AAA).

### In Utero Electroporation

Sprague Dawley (SD) rats were used for *in utero* electroporation experiment as described previously (*29, 30*). Briefly, electroporation was performed at E15-16. Pregnant rats were anesthetized with chloral hydrate and embryos were exposed in the uterus. Saline solution containing 0.05% Fast Green FCF dye (Sigma) and expression plasmids pAAV-EGFP-ASIC1a, pAAV-mCherry-Lact-C2, or control vector pAAV-GFP/ mCherry under the control of a human Synapsin I promoter was injected (1 μg/μl) into the lateral ventricle of embryonic brains. The embryo’s cerebral wall was electroporated with five square electrical pulses (50 V, 50 ms duration at 100 ms intervals) generated by an ECM-830 BTX Electro Square Porator (Harvard Apparatus Inc.). The pups were sacrificed at P18. Brains were processed and sections were stained with anti-GFP and anti-DsRed antibodies to visualize neuronal morphology and protein locations (see below *Immunocytochemistry*).

### GST Fusion Protein and Phospholipid Binding Assay

To express GST fusion proteins, the intracellular N- and C-termini of human ASIC1a (Nt_hASIC1a_: aa1-41, Ct_hASIC1a_: aa465-528) and hASIC2a (Nt_hASIC2a_: aa1-40, Ct_hASIC2a_: aa462-512) were amplified by PCR and inserted into pGEX-4T1 vector between the BamHI and EcoRI sites. To purify GST fusion proteins, the constructs were transformed into *E.coli* BL21 (DE3) pLysS competent cells (Tiangen, China). After growth to OD_600_ = 1 in a SuperBroth medium supplemented with ampicillin, the expression of GST fusion proteins was induced with isopropylthiogalactoside (IPTG, Sigma) at 16 °C overnight. Then, *E.coli* cells were lysed by using BugBuster Kit (Novagen) and purified with Glutathioine Sepharose 4B (GE healthcare) according to the manufacturer’s protocol. The expression levels and molecular weights of GST fusion proteins were checked by Coomassie blue staining, while an empty pGEX vector was used as a control. The concentration of eluted GST fusion protein was determined by absorbance at 280 nm.

For lipid strip-binding assay, PIP strips (Echelon) pre-blocked with bovine serum albumin (BSA) were incubated with 1 μg/ml of purified GST fusion proteins for 1 hr at room temperature and then washed 3 times with phosphate-buffered saline (PBS) or Tris-buffered saline (TBS) containing 0.1% Tween 20 (PBS-T or TBS-T, respectively), followed by incubation with anti-GST mAb (1:2000, Sigma) and anti-mouse IgG-HRP (1:2000, Sigma) sequentially. Bound proteins were detected with the ECL detection system (Thermo Fisher Scientific).

For lipid bead-binding assay, purified GST fusion proteins (5 μg) were diluted in 500 μl binding buffer (50 mM Tris, 150 mM NaCl, 0.25 % NP-40, pH 7.5) and incubated with 30 μl of PS-conjugated Sepharose beads (Echelon) for 1 hr. After three quick washes with the binding buffer, bound proteins were eluted from the beads by boiling with the SDS sample buffer for 10 min, and then detected by immunoblotting.

### Cell Culture and Transfection

Primary cortical neurons from SD rats (E18) and wild-type or *Asic1a^-/-^* mice (P0) were dissociated and maintained using a standard protocol (*31*). Briefly, cerebral cortices of the rodents were dissected in D-Hank’s solution, dissociated in 0.05% trypsin, and then plated on poly-D-lysine (Sigma) coated cover glasses or culture dishes (Corning) (2×10^5^ cells/ml for electrophysiology and immunocytochemistry; 1×10^6^ cells/ml for biochemistry). The plating medium was DMEM (Gibco) with 10% fetal bovine serum (FBS), 10% F12 and 1% Glutamax (Gibco), a half of which was replaced by Neurobasal medium (Gibco) containing 2% B27 nutrient mixture (Gibco) and 1% Glutamax one day after cell plating. Chinese hamster ovary (CHO) cells were grown in F12 medium with 10% FBS and 1% Glutamax.

All cells were cultured under 37 °C and 5% CO_2_ humidified conditions. Transient transfection was performed with the use of HilyMax liposome transfection reagent (Dojindo Laboratories) or CalPhos Mammalian Transfection Kit (Clontech). Unless otherwise noted, all media and supplements were purchased from Gibco, Life Technologies Co., and equal amounts of plasmids were used for every combination of constructs for co-transfection. Protein extraction, electrophysiology and immunostaining were performed at least 24 hr after transfection.

### Phospholipid Uptake

1-palmitoyl-2-(dipyrrometheneboron difluoride) undecanoyl-sn-glycero-3-phospho-L-serine (TopFluor-PS) was purchased from Avanti Polar Lipids. Lipid dissolved in chloroform was aliquoted and dried under nitrogen gas in a glass tube, and then resuspended in methanol. Neuronal cultures of 5-6 weeks old were incubated with 1 μM TopFluor-PS in extracellular solution (ECS, 150 mM NaCl, 5 mM KCl, 1 mM MgCl_2_, 2 mM CaCl_2_, 10 mM glucose, 10 mM HEPES, pH 7.4) for 10 min on ice, followed by 10 min backwash with ice-cold ECS containing 4% (wt/vol) fatty acid-free BSA before the original culture medium was added back. After incubation for 3-4 hr at 37 °C, cells were fixed and imaged by confocal microscopy. To visualize dendritic morphology of individual neurons, pAAV-mCherry was transfected into the cortical neurons at days *in vitro* (DIV) 7.

### Annexin V Binding Assay

Annexin V-FITC Apoptosis Detection Kit (Sigma-Aldrich) was used for detecting surface PS-exposure, according to the manufacturer’s instructions. Briefly, CHO cells were plated at 1 x 10^6^ cells/well in a 6-well plate and cultured for 24 hr in 2 ml of F12 medium (Gibco) supplemented with 10% FBS and 1% Glutamax (Gibco) (5% CO_2_ and 37°C). Staurosporine (1 μg/ml) was applied for 2 hr to induce apoptosis and PS externalization, followed by washing gently with PBS (*32*). Then, 500 μl of the binding buffer (10 mM HEPES/NaOH, pH7.5, 140 mM NaCl, 2.5 mM CaCl_2_) containing 5 μl of Annexin V-FITC conjugates (50 μg/ml) was added to each well and the cultures were incubated for 10 min at room temperature. After the incubation, cells were washed with the binding buffer, fixed with 4% paraformaldehyde for 15 min, and then washed and immersed in the binding buffer again for imaging. Images were acquired from five random microscopic fields of each well using an Olympus IX71 inverted fluorescent microscope. Numbers of total cells (bright field) and Annexin V-positive cells were counted, and FITC intensities were compared between control and shRNA-transfected groups (DsRed-positive). Experiments were performed in triplicates.

### Antibody Feeding Assay

Cultured neurons were transfected with wild-type ectodomain HA-tagged ASIC1a (HA-ASIC1a) using the calcium phosphate precipitation transfection method at DIV7. After 48 hr, cells were blocked (5% FBS in PBS) for 15 min, followed by incubation with polyclonal anti-HA antibodies for 15 min at 4°C. After removing the unbound antibody by washing three times with PBS at 4°C, the cells were treated with the desired drugs or peptides for varying time periods, followed by incubation in conditioned medium to allow internalization for additional 10 min at 37°C. Then, the cells were fixed in 4% paraformaldehydein PBS for 15 min and incubated with a saturable amount of Alexa 568-conjuagted secondary antibodies to detect surface HA-ASIC1a. After that, the cells were permeabilized with 0.1% Triton X-100 and then incubated with Alexa 647-conjugated secondary antibodies to detect internalized HA-ASIC1a. The internalization ratio of HA-ASIC1a was calculated by dividing the endocytic (Alexa 647) to the surface (Alexa 568) fluorescent signals of individual cells.

### Surface Biotinylation Assay

Surface biotinylation assay was performed following the established protocol (*33*). In short, transfected cells were washed three times with ice-cold PBS (with 1 mM MgCl_2_ and 2.5 mM CaCl_2_), and then 0.25 mg/ml sulfo-NHS-LC-biotin (Thermo Fisher Scientific) was applied to the cultures followed by incubation for 30 min at 4°C. After the incubation, cells were washed three times with 100 mM glycine/PBS to quench the excess biotin, followed by three additional washes with PBS and then homogenization in the lysis buffer (20 mM Tris-HCl, 137 mM NaCl, 10% glycerol, 1% NP-40, 2 mM EDTA, pH 8). After excluding the nuclei by centrifugation, the lysates were incubated and rotated with immobilized NeutrAvidin agarose resins (Pierce) for more than 3 hr at 4°C. The beads were washed with 1% Triton X-100/PBS and then boiled in SDS loading buffer to elute the bounded proteins, which were subsequently subject to immunoblotting together with samples of total proteins (input). Image J 1.52d (NIH, USA) was used to quantify the band intensities of immunoblots.

### Immunocytochemistry

At 24 hr after transfection, cells cultured on glass coverslips were fixed for 15 min with 4% paraformaldehyde/PBS at 37 °C and blocked with 10% FBS/PBS for one hr at room temperature. Triton X-100 (0.1%) was added into the blocking buffer to permeabilize the cells when needed. Cells were incubated overnight with anti-GFP (1:500, Invitrogen), anti-DsRed (1:500, Clontech), anti-Synapsin I (1:1000, kind gift from Dr. Nan-Jie Xu, Shanghai Jian Tong University School of Medicine), anti-PSD-95 (1:500, Abcam), anti-cMyc mAb (1:1000), anti-HA mAb (1:1000, SIGMA), anti-HA pAb (1:500, GeneTex), or anti-EEA1 (1:100, Cell Signaling Technology) at 4°C, followed by rinsing with PBS to remove unbound antibodies. Then, the cells were incubated with AlexaFluor 488-, 543- or 633-conjugated secondary IgG (1:2000, Invitrogen) for 2 hr at room temperature, rinsed with PBS and afterwards mounted with the fluorescent mounting medium (Dako).

Immunofluorescent images were obtained using either a Nikon A1R confocal laser scanning microscope (Nikon) equipped with Plan Apo VC 20x DIC N2 (NA 0.75) dry objective /Apo 60x Oil λS DIC N2 (NA 1.40) oil-immersion objective lens, and analyzed in Nikon NIS-Elements AR 4.40; or a Leica TCS SP8 confocal laser scanning microscope (Leica Microsystems) with Leica HC PL APO CS2 40x (NA 1.30) /63x (NA 1.40) oil-immersion objectives, and analyzed using Leica LAS X software (Leica Microsystems). In individual experiments, cells from the same culture and transfection preparation were treated with vehicle or drugs and compared with each other, while the images for all conditions were obtained on the same equipment with identical acquisition parameters (for example, laser power, gain, offset, pinhole and scanning speed, etc.) and quantified using Image J 1.52d software (National Institutes of Health) with identical parameters. To obtain the entire morphology of a cultured neuron or a dendrite, a z-stack (0.5-1 μm each step) of optical sections was captured.

### Super-resolution imaging

For super-resolution imaging, structured illumination microscopy (SIM) images were acquired using a Nikon Structured Illumination super-resolution microscope (N-SIM) with a 100x oil-immersion objective (1.4 NA). SIM imaging was performed at Institute of Brain Science, Fudan University, and the National Center for Protein Science Shanghai. Z-stack (0.2 μm each step) images of dendritic spines were processed and reconstructed into three-dimensional structures using Nikon Elements software. In order to verify the results from Nikon N-SIM images, we also used Zeiss LSM 800 microscope with an Achroplan 100x (1.4 NA) oil-immersion objective and Airyscan array detector to obtain the super-resolution images and analyzed the images using the ZEN Microscope Software.

### Image Analysis

Image analysis was performed using NIS-Elements AR or Image J software. To measure the areas of different compartments of the cortical neurons, individual neurons or dendrites were converted to binary masks using intensity threshold. ROIs for various compartments were outlined manually from the original threshold mask in each channel. The mean intensity or integrative density for individual ROIs in each channel was measured automatically. To quantify the degree of signal colocalization between different channels, Pearson’s correlation coefficient was employed. Line scans were performed on the maximum intensity z projection images for dendritic spine analysis. Only spines on secondary and tertiary dendrites of cortical neurons were analyzed. Spine types were classified based on the morphology and traditional classification method (*34*). Sample groups for direct comparison were treated simultaneously and imaged with the same parameters. For each group, at least 5 neurons and 100 spines were analyzed. Three-dimensional shaded volume rendering of dendritic spines was created in NIS-Elements AR.

### Fluorescent recovery after photobleaching (FRAP)

FRAP experiments were performed as previously reported (*18, 28*). Briefly, rat cortical neurons were transfected with cDNA coding for EGFP or EGFP-ASIC1a proteins at DIV7 and transferred to the imaging chamber at DIV21. To assess ASIC1a dynamics, images were acquired every 500 ms. After a short baseline recording (20 s - 1 min), fluorescent signal of EGFP in a small defined region of interest (ROI) was eliminated by a bleaching laser train (2-s pulse duration, 5 scans, 25% power) from a 488-nm argon (100 mW, Melles Griot) or solid-state (30 mW, Pavilion) laser source. The bleached ROIs (< 1 μm^2^) were outlined at the middle of spine heads and dendritic shafts on the tertiary branches of the dendrites of the cortical neurons. The amount of recovery at each time point was calculated as FRAP_t_ = (F_t_ - F_0_)/(Fi - F_0_), where F_t_ is the fluorescence in the bleached area at time t, F_0_ is the fluorescence of the same field immediately after bleaching, and F_i_ is the initial intensity of the field before photobleaching.

### Topology Prediction

Server-based online programs including TMHMM2.0, TMPred, HMMTOP, CCTOP, TopPred, PredictProtein, ConPredII and TopGraph were used to depict the secondary structure, hydrophobicity, and topology of ASIC1a proteins and profile the regions of second transmembrane domain (TMII) and the cytoplasmic C-terminus of ASIC1a. Lipid-cytoplasmic interface of ASIC1a was also evaluated by helicity index generated using PSIPRED. Gene sequence alignment was performed using AlignX from the Vector NTI software package (Thermo Fisher Scientific).

### Whole-cell Patch-clamp Recordings

Solutions and methodology for electrophysiological recording of acid-induced currents were as previously described from our laboratory (*35*). In brief, all chemicals for whole-cell recordings were purchased from Sigma. The extracellular solutions containing 150 mM NaCl, 5 mM KCl, 1 mM MgCl_2_, 2 mM CaCl_2_, and 10 mM glucose were adjusted to various pH values with 10 mM HEPES (pH 6.0–7.4) or MES (pH < 6.0). Patch pipettes (3-5 MΩ) were filled with the standard pipette filling solution which contained 120 mM KCl, 30 mM NaCl, 0.5 mM CaCl_2_, 1 mM MgCl_2_, 5 mM EGTA, 2 mM Mg-ATP, and 10 mM HEPES (pH 7.2). Whole-cell voltage clamp recordings were performed at room temperature (22-25°C) with an Axopatch 700B amplifier and Digidata 1320A interface (Axon Instruments). Under voltage clamp conditions, the membrane potential was held at −60 mV throughout the experiment and compensation for capacitance and series resistance was manually applied in most experiments. Data were sampled and analyzed with pClamp 10 software (Axon Instruments).

### Statistical Analysis

Data analysis was performed using OriginPro 8 SR0 8.0724 (OriginLab) and GraphPad Prism 6.01 (GraphPad Software) software. Data are presented as mean ± SE (s.e.m). Statistical comparisons between two groups were performed by unpaired or paired Student’s *t*-tests. One-way ANOVA (either parametric or nonparametric) followed by *post hoc* tests as described in each figure legend was used for the comparison of multiple groups. *p* values < 0.05 are considered statistically significant. For *p* values less than 0.0001, we provide a range rather than the exact number.

## Results

### PS interacts with ASIC1a channel

To find if ASIC1a binds to any phospholipid species, we screened the main phospholipids immobilized on nitrocellulose membranes (*36*) for their ability to retain glutathione S-transferase (GST)-tagged cytoplasmic amino- and carboxyl-termini of human ASIC1a, GST-Nt_ASIC1a_ and GST-Ct_ASIC1a_, respectively **(Fig. 1A)**. GST fusion proteins of full-length ASIC1a were not soluble when expressed in *E.coli* (data not shown) and therefore could not be tested. We found that GST-Ct_ASIC1a_, but not GST alone or GST-Nt_ASIC1a_, specifically bound to two acidic phospholipid species, phosphatidylserine (PS) and phosphatidic acid (PA), without interacting with phosphoinositides (PIPs) or other phospholipids **(Fig. 1A)**. In both TBS (Tris-based) and PBS (phosphate-based) buffering systems, this assay consistently showed the same binding pattern, with a higher affinity of PS to ASIC1a than PA (**fig. S1, A and B**), distinct from the lipid-binding profiles of other membrane receptors (**fig. S1, C to E**). We further validated the interaction between PS and ASIC1a C-terminus by using PS-conjugated agarose beads to pull down GST-tagged proteins (**Fig. 1B**), and found that the PS-beads, but not control lipid beads, successfully pulled down GST-Ct_ASIC1a_ more efficiently than GST alone or GST-Nt_ASIC1a_.

**Figure 1.**
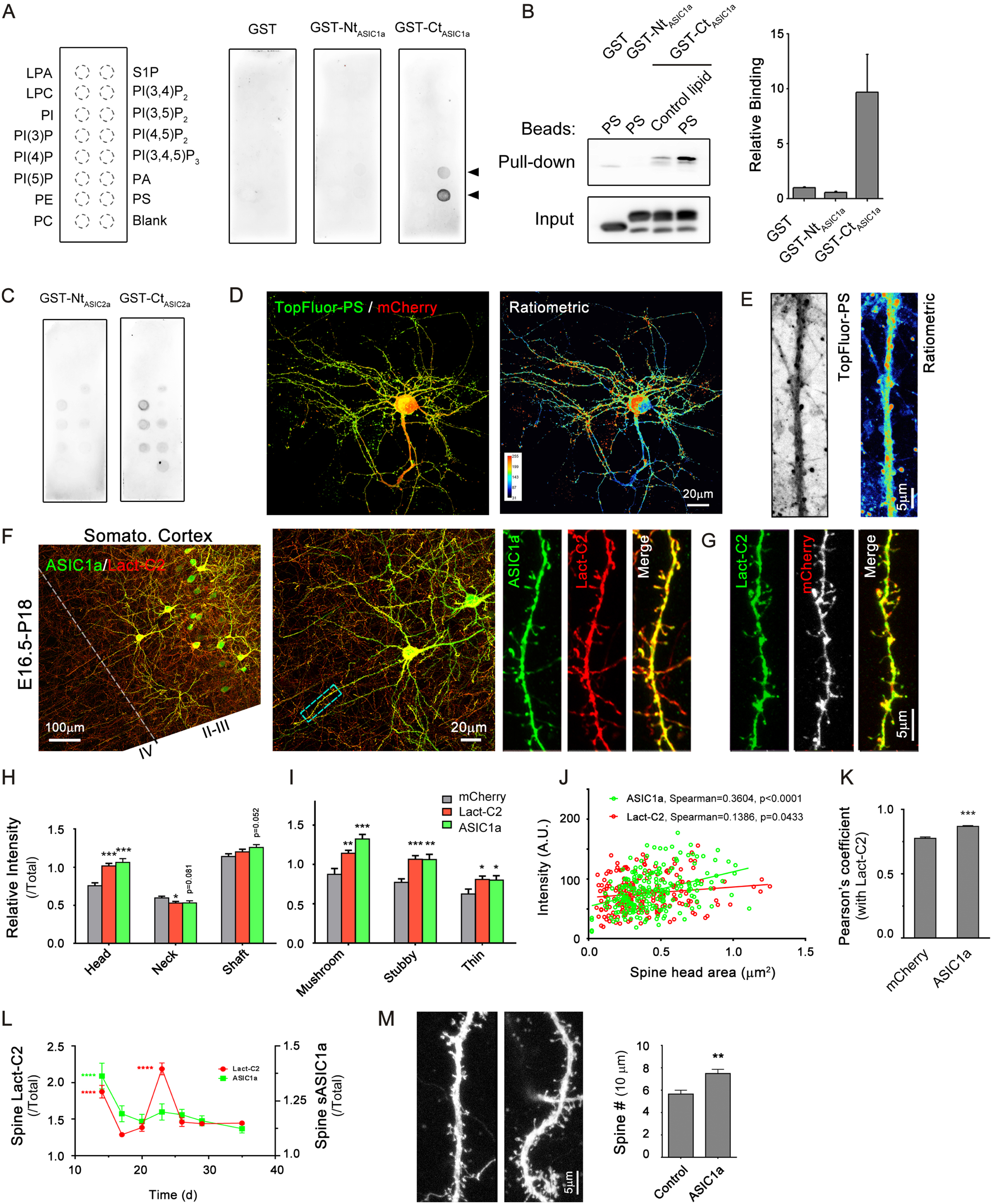
ASIC1a preferentially binds to anionic phosphatidylserine. **A.** Lipid-protein overlays. The nitrocellulose membrane strip was spotted with 15 different kinds of lipid molecules (100 pmol each; left diagram). Purified GST fusion proteins: GST alone (left), GST-Nt_ASIC1a_ (middle), and GST-Ct_ASIC1a_ (right), were incubated with the strip in TBS buffering system and then detected with anti-GST antibodies. Arrowheads, positive GST signals. LPA, lysophosphatidic acid; LPC, lysophosphatidylcholine; PI, phosphatidylinositol; PE, Phosphatidylethanolamine; PC, phosphatidylcholine; S1P, sphingosine-1-phosphate; PA, phosphatidic acid; PS, phosphatidylserine. **B.** Lipid-protein pull-down assay. GST fusion proteins as in (A) were pull-downed with lipid-coated agarose beads in HEPES buffering system (mean lipid concentration = 10 μM), and probed with anti-GST antibodies. Left, representative immunoblots of pull-down fraction and input of GST-fusion proteins: GST (∼27 kDa), GST-Nt_ASIC1a_ (∼35 kDa) and GST-Ct_ASIC1a_ (∼32 kDa); Right, statistic results of pull-down assay normalized to intensity values of control group by GST. Data are mean ± SEM. **C.** Strip screening for ASIC2a. GST fused N- and C-termini of ASIC2a: GST-Nt_ASIC2a_ (left) and GST-Ct_ASIC2a_ (right), respectively, were detected as in (A). **D.** Representative confocal images (left) of a DIV28 rat cortical neuron transfected with mCherry at DIV7. TopFluor-PS (1 µM) was applied for 10 min on ice. After 10 min wash on ice and 3-4 hr redistribution process at 37 °C, cells were fixed and directly imaged under confocal microscope. Ratiometric image for TopFluor-PS over mCherry of the same view is also shown (right). **E.** Representative confocal images of a neuronal dendrite from samples processed the same way as in (D). Scale bar, 5 μm. **F.** Representative confocal images of coronal slice (left) of a P18 rat brain that was transfected with EGFP-ASIC1a and mCherry-Lact-C2 by *in utero* electroporation at E16.5. Brain slices were immunostained by antibodies of the corresponding fluorescent proteins. High-magnification example images of layer II/III pyramidal neurons (middle) and a dendrite (right) in the same view are also shown. **G.** Representative images of another neuronal dendrite from rats transfected with EGFP-Lact-C2 and mCherry and detected as in (F). **H & I.** Quantification of distributions of indicated proteins in different compartments of neuronal dendrites (H) or different types of dendritic spines (I) for neurons processed as in (F) and (G). Mean fluorescent intensities of indicated proteins in each dendritic area (relative to the whole neuronal area) are compared with the control group that expressed mCherry. N = 30-60, \**p* < 0.05, \*\**p* < 0.005, \*\*\**p* < 0.001 (one-way ANOVA with Fisher’s least significant difference (LSD) post hoc). **J.** Correlation plots of signal intensities of ASIC1a (green, R^2^ = 0.1441, slope = 57.92 ± 9.474, *p* < 0.0001, F-test, n = 224 spines) and Lact-C2 (red, R^2^ = 0.0200, slope = 17.64 ± 8.503, *p* = 0.0392, F-test, n = 213 spines) against spine head areas, n = 11 cells. **K.** Quantitative colocalization analysis for mCherry or ASIC1a with Lact-C2. Pearson’s correlation coefficients were obtained by analysis of 130 and 206 spines from at least 11 cells. \*\*\**p* < 0.001 (Student’s *t*-test). Error bars indicate SEM. **L.** Spine levels of ASIC1a and Lact-C2 during neuronal development *in vitro* (2-5 weeks). For DIV14 and DIV23, \*\*\*\**p* < 0.0001 (Kruskal-Wallis’s test), n > 70. **M.** Spine density (numbers / 10 µm) of ASIC1a-transfected hippocampus neurons in cultured organotypic brain slices. \*\**p* < 0.01 (Student’s *t*-test), n > 16 neurons, dendritic lengths analyzed: Control, 4048 μm; ASIC1a, 7575 μm.

To test whether this phospholipid-binding pattern is conserved in the ASIC family, we examined phospholipid binding of cytoplasmic N- and C-termini of ASIC2a. Although both GST-Nt_ASIC2a_ and GST-Ct_ASIC2a_ showed some weak binding to various phospholipid species, with GST-Ct_ASIC2a_ being stronger than to GST-Nt_ASIC2a_, they did not exhibit a specific interaction with PS (**Fig. 1C and fig. S1B**). Therefore, the high affinity PS binding is ASIC1a specific and ASIC1a C-terminus is responsible for this interaction.

### Spine enrichment of PS and ASIC1a

Previous studies showed that PS has asymmetric cellular distribution in yeast (*Saccharomyces cerevisiae*) (*37*). The distribution pattern of PS in mammalian neurons should also be of great interest because neurons usually have the most delicate morphological structures and undergo dynamic changes even in mature stages. ASIC1a is highly expressed in the dendritic spines of neurons and functions as a synaptic pH-sensor (*18*). The finding that ASIC1a interacts with PS prompted us to ask whether PS is also enriched in the dendritic spines. To test this, we first used fluorescent TopFluor-PS, a synthetic PS analogue, to examine PS distribution and dynamics in cultured rat cortical neurons (**Fig. 1, D and E**) (*38*). Once applied to the culture, TopFluor-PS was taken up rapidly by the neurons and concentrated in endocytic organelles, showing bright and round vesicular structures. After 3-4 hr incubation at 37 °C, the fluorescent signals of TopFluor-PS were redistributed and largely associated with the PM. Notably, compared to the fluorescent signals of soluble mCherry control (transfected at DIV7), TopFluor-PS was more enriched at the head areas of dendritic spines (**Fig. 1, D and E**). To examine the *in vivo* localizations of PS and ASIC1a in cortical neurons, we employed a genetically encoded PS probe: C2 domain of lactadherin (Lact-C2) (*8*). We fused the open reading frame of Lact-C2 with either mCherry or EGFP, and co-transfected either EGFP-ASIC1a with mCherry-Lact-C2 or EGFP-Lact-C2 with the mCherry vector into the radial glial progenitors in the cerebral cortex of E15.5 rats using *in utero* electroporation (IUE) (**Fig. 1, F to K**). Interestingly, confocal images of brain sections of transfected P18 rats consistently showed higher enrichment of EGFP-Lact-C2 than soluble mCherry in spine head areas of cortical neuron dendrites *in vivo* (**Fig. 1, G and H**), consistent with the finding made using TopFluor-PS in cultured neurons. The enrichment of Lact-C2 was found in mushroom, stubby, and thin spines, categorized according to their morphology and location, and more so in the mushroom and stubby spines than in thin spines (**Fig. 1I**), showing a weak correlation with the spine head area (**Fig. 1J**). More importantly, clear overlaps between Lact-C2 and ASIC1a signals were found in the spines (**Fig. 1F**), showing significantly higher Pearson’s coefficient than that between Lact-C2 and soluble mCherry alone (**Fig. 1K**). These indicate a close colocalization between ASIC1a and PS in dendritic spines.

Notably, the spine enrichment of PS during early development was not constant (**Fig. 1L**). In two time periods, which represent the stages of spine growth (about 2 weeks) and branching (about 3-4 weeks), Lact-C2 in dendritic spines had nearly twice the intensity as compared to other periods (**Fig. 1L**), indicating that PS recruitment to dendritic spines is dynamically regulated. Interestingly, the spine expression of ASIC1a was highest at 2 weeks and then maintained at a lower level at P17-P35 (**Fig. 1L**). Also, overexpression of ASIC1a significantly increased dendritic spine density of hippocampal neurons in organotypic brain slices (**Fig. 1M**) (*19*). Therefore, not only PS (*5*), but ASIC1a also contributes to spine development.

To further confirm the spine localization of PS and ASIC1a, we examined the expression patterns of Lact-C2 and ASIC1a in cultured rat cortical neurons (**fig. S2A**). Compared with the relatively even distribution of mCherry (*right*), both Lact-C2 (*left*) and ASIC1a (*middle*) showed higher expression levels in dendritic spines as compared to the shaft. By contrast, the Lact-C2 mutant, Lact-C2-W26A/W33A/F34A (Lact-C2-AAA), which has the three key PS-binding residues mutated to alanine (*5*), exhibited the relatively even distribution in soma and dendritic shafts as EGFP or mCherry, showing markedly reduced expression in dendritic spines compared to the wild type Lact-C2 (**fig. S2, B to D**). Additionally, we showed that Lact-C2 and ASIC1a colocalized substantially with postsynaptic marker, PSD-95, presynaptic marker, Synapsin I, and Phalloidin (for F-actin) in rat cortical neurons at DIV 28 (**Fig. 2, A to C**), further demonstrating that PS and ASIC1a are both highly enriched in dendritic spines. The specificities of synaptic markers were validated in control experiments (**fig. S2E**). Taken together, the results from both *in vivo and in vitro* assays suggest that the anionic phospholipid, PS, directly binds with the cytoplasmic C-terminus of ASIC1a, and this may be important for the preferential enrichment of ASIC1a in dendritic spines.

**Figure 2.**
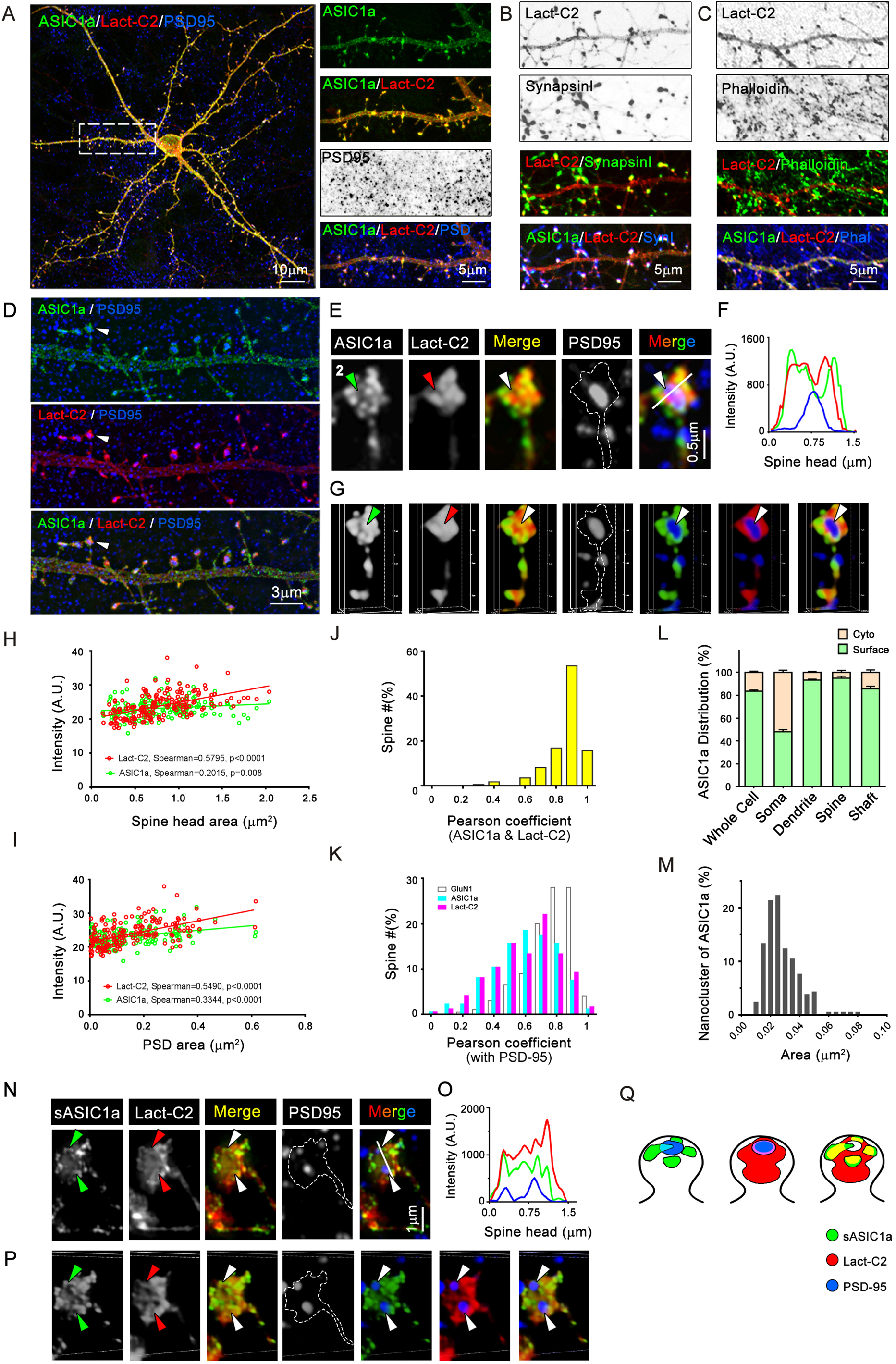
Synaptic localizations of PS and ASIC1a. **A-C.** Representative confocal images of a DIV28 rat cortical neuron that was transfected with EGFP-ASIC1a and mCherry-Lact-C2 on DIV7. After fixation, primary cultures were immunostained with the antibodies for PSD-95 (A), Synapsin I (B), and corresponding fluorescent proteins. F-actin was visualized by staining with Alexa Fluor 647-phalloidin (C). **D.** Representative super-resolution images of a DIV28 rat cortical neuron that was transfected with EGFP-ASIC1a and mCherry-Lact-C2 on DIV7, imaged with structured illumination microscopy (SIM). After fixation, primary cultures were immunostained with the antibodies for PSD-95 and corresponding fluorescent proteins. Arrowheads indicate the spine shown in E. More details are shown in Supplementary Figure 3. **E.** SIM z stack maximum projection images of a representative dendritic spine in (D). Arrowheads, PSD area; white line, line scan area in (F). **F.** Intensity profiling of line scan through the spine head in (E). ASIC1a, green line; Lact-C2, red line; PSD-95, blue line. **G.** 3D reconstruction of the same spine in (E). Arrowheads, PSD area. **H.** Correlation plots of signal intensities of spine ASIC1a (green, R^2^ = 0.0254, slope = 1.036 ± 0.4917, *p* = 0.0367, F-test) and Lact-C2 (red, R^2^ = 0.2576, slope = 4.672 ± 0.6082, *p* < 0.0001, F-test) against spine head areas, n = 172 spines from 7 cells. **I.** Correlation plots of signal intensities of spine ASIC1a (green, R^2^ = 0.1159, slope = 7.001 ± 1.483, *p* < 0.0001, F-test) and Lact-C2 (red, R^2^ = 0.2811, slope = 15.45 ± 1.895, *p* < 0.0001, F-test) against PSD areas, n = 172 spines from 7 cells. **J.** Frequency distribution of Pearson’s correlation coefficients between ASIC1a and Lact-C2 for 172 dendritic spines from 7 cells. **K.** Frequency distributions of Pearson’s correlation coefficients between PSD-95 and GluN1 (open, 200 spines from 10 cells), ASIC1a (cyan, 172 spines from 7 cells) or Lact-C2 (pink, 172 spines from 7 cells). \*\*\**p* < 0.001 between ASIC1a/Lact-C2 group and GluN1 group; no significance between ASIC1a and Lact-C2 groups (one-way ANOVA with Tukey’s post hoc). **L.** Percentage of surface and cytoplasmic ASIC1a in different compartments of rat cortical neurons at 5-6 weeks. The values are: surface ASIC1a, 83.6% ± 1.0% (whole cell), 48.0% ± 1.9% (soma, \*\*\*\**p* < 0.0001, vs. whole cell), 93.3% ± 0.8% (dendrite, \*\*\*\**p* < 0.0001, vs. whole cell), 95.0% ± 1.7% (spine head, \*\*\**p* < 0.001, vs. whole cell), and 85.7% ± 2.2% (dendritic shaft, no significance, vs. whole cell), n = 200 spines from 11 cells, one-way ANOVA with Tukey’s post hoc analysis. **M.** Frequency distribution of sizes of surface ASIC1a nanoclusters. **N.** SIM z stack maximum projection images of a dendritic spine of a DIV28 mouse *Asic1a*^-/-^ cortical neuron transfected with HA-ASIC1a and mCherry-Lact-C2 on DIV7 and stained for surface HA, total mCherry, and total PSD-95. Arrowheads, PSD area; white line, line scan area in (O). **O.** Intensity profiling of line scan through the spine head in (N). Surface ASIC1a, green line; Lact-C2, red line; PSD-95, blue line. **P.** 3D reconstruction of the same spine in (N). Arrowheads, PSD area. **Q.** Schematic diagram for synaptic localizations of different proteins. Green, surface ASIC1a (left); blue, PSD-95; red, Lact-C2; cyan (left), pink (middle), or yellow (right), colocalization signal.

### Synaptic localizations of ASIC1a and PS

Spines and synaptic sites are finely organized structures that contain distinct microdomains both spatially and functionally (*39*). To understand the distributions of ASIC1a and PS at the spine head areas and, particularly, their relationships in nanoscales, we took the advantage of the robust imaging power of structured illumination microscopy (SIM) technique. In cultured cortical neurons co-transfected with EGFP-ASIC1a and mCherry-Lact-C2, and immunostained with the antibodies for EGFP, mCherry, and PSD-95, super-resolution SIM images revealed that ASIC1a existed as nanometer-sized clusters while Lact-C2 had a slightly smoother appearance than ASIC1a in spine heads (**Fig. 2, D to G, and fig. S3, A to D**). Both proteins consistently showed higher levels at the spine head areas than in spine neck or dendritic shaft. In the spine head, they spread around the entire area such that the areas occupied by ASIC1a or Lact-C2 strongly correlated with the sizes of the spine heads, as well as the postsynaptic (PSD) areas (indicated by staining with PSD-95) (f**ig. S3, E to G**). The signal intensities of ASIC1a and Lact-C2 in individual spine heads also positively correlated with the spine sizes and PSD areas (**Fig. 2, H and I**). These data indicate that ASIC1a and PS are present throughout the entire spine head, but the levels may differ according to spine maturation levels. Notably, even at this high resolution, the signals of ASIC1a and Lact-C2 are still well colocalized. The Pearson’s correlation coefficient between the two proteins exceeded 0.8 in > 86% of the spines (**Fig. 2J**).

To our surprise, a closer inspection of the super-resolution structures of the spine heads revealed that the spine-enriched ASIC1a and Lact-C2 frequently exhibited peri-synaptic localizations that are adjacent to or partially overlapping with PSD-95 **(Fig. 2, D to G)**. Line scans through the spine heads on the Z-max-projection images showed that the peak of PSD-95 did not overlap with ASIC1a or Lact-C2 (**Fig. 2, E and F, and fig. S3, B and C**). In 3D reconstructions, almost all the dendritic spines had the stereoscopic PSD-95 fluorescent signals surrounded by, rather than co-labeled with, ASIC1a or Lact-C2 signals **(Fig. 2G and fig. S3D)**. Compared to SEP-GluN1, a control of postsynaptic receptors that are highly overlapped with PSD-95 **(fig. S4)**, the colocalization of ASIC1a or Lact-C2 with PSD-95 was considerably weaker (**Fig. 2K**). These data suggest that a large portion of ASIC1a and PS localize at peri-synaptic regions, rather than at the core of PSD. Moreover, we also employed Zeiss LSM800 confocal microscopy with Airyscan array detector, which captures super-resolution images via a distinct mechanism from SIM, to verify our results **(fig. S5)**. Because LSM800 microscopy can freely change the imaging detectors between the normal one and Airyscan detector array to capture the same field-of-view, we were able to compare the confocal images with Airyscan images of the same samples. The nanoscale localizations of ASIC1a and Lact-C2 revealed by Airyscan images confirmed that the spine-enriched ASIC1a and Lact-C2 were highly colocalized at the peri-synaptic sites of dendritic spine heads **(fig. S5)**.

Given that only the surface-expressed ASIC1a (surface ASIC1a) is functionally relevant in terms of sensing extracellular acidification, we next determined the synaptic localization of surface ASIC1a. First, we found that the vast majority of ASIC1a proteins is present on the cell surface (83.6% ± 3.5%), especially in dendrites (93.3% ± 2.6%) and spines (95.0% ± 5.4%) (**Fig. 2L and fig. S6**). Next, we examined the relationships between surface ASIC1a and Lact-C2 or PSD-95 using a cDNA construct encoding ASIC1a with a hemagglutinin (HA) tag inserted in the ectodomain (HA-ASIC1a) as previously described (*18, 27, 28*). Expression of HA-ASIC1a in cultured *Asic1a^-/-^* mouse cortical neurons allowed distinction and quantification of surface and total ASIC1a proteins by performing immunostaining with anti-HA under non-permeabilized and permeabilized conditions, respectively. When imaged by SIM, surface HA-ASIC1a clusters were found throughout the dendrites and spines, with the mean densities of 22.16 ± 0.90, 16.14 ± 1.35, 18.73 ± 0.99 per μm^2^ on spine head, spine neck and dendritic shaft, respectively (**fig. S3, H to K**). The mean diameter of the surface HA-ASIC1a clusters was 0.159 ± 0.048 μm (**fig. S3L**) and the mean size was 0.027 ± 0.011 μm^2^ (**Fig. 2M**). The line scan result on Z-max-projection images and 3D reconstruction also showed similar results as that of the total ASIC1a (EGFP-ASIC1a) (**Fig. 2, N to Q**).

Together, with much higher spatial resolutions than Annexin V-GFP labeling results (*40*), our data provide more precise information on the distribution of PS and ASIC1a in cortical neuron dendrites. Although controversy exists regarding ASIC1a subcellular distributions in neurons (*19, 41*), our data clearly show that ASIC1a and Lact-C2 are enriched and well-colocalized at peri-synaptic sites in spine heads, and the expression of ASIC1a and PS there positively correlates with the sizes of the spine heads and PSD. Since both PS (*42, 43*) and ASIC1a (*44, 45*) exist in high levels and undergo regulated processing in neurons, our data also indicate that ASIC1a and PS may play important roles in spine morphology and synaptic function.

### PS promotes surface expression of ASIC1a

To define the functional role of PS on ASIC1a channels, we employed several approaches to manipulate intracellular PS levels. PS synthase I (PtdSSI, PSSI) and PS synthase II (PtdSSII, PSSII) are the two major PS synthases using different substrates: phosphatidylcholine (PC) and phosphatidylethanolamine (PE), respectively (**Fig. 3A**) (*46, 47*). To decrease the cellular level of PS, we designed four shRNA candidates targeting PSSI or PSSII to individually knockdown these lipid biosynthetic enzymes (**Fig. 3, B and C**). The myc-tagged mouse PSSI or PSSII was co-transfected with each candidate shRNA construct into CHO cells and the knockdown efficiencies were examined by immunoblotting with anti-myc antibody (*48*). All of the four PSSI shRNAs and two of the four PSSII shRNAs resulted in significant knockdown. The shRNA candidates with the highest knockdown efficiency and sequence conservation across species (i.e., shPSSI-#456 and shPSSII-#600) were chosen to transfect cortical neurons to target endogenous PSSI and PSSII *in vitro* (**Fig. 3D**). We found the surface level of ASIC1a to be markedly decreased in the shPSSI-transfected group and the shPSSI and shPSSII co-transfected group, but not the shPSSII-transfected group, indicating that PSSI plays a role in promoting the surface level of ASIC1a.

**Figure 3.**
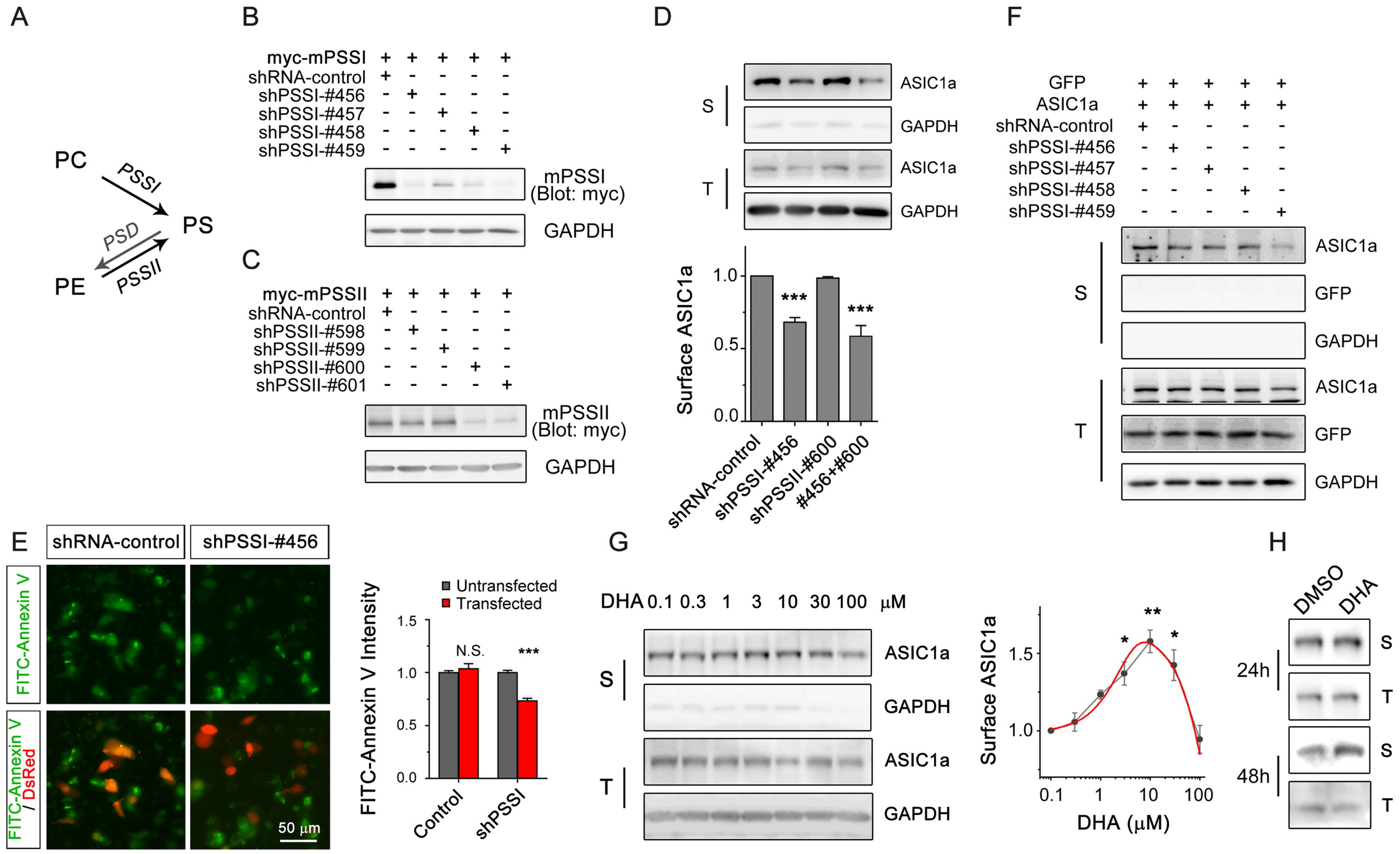
PS regulates surface expression of ASIC1a. **A.** Biosynthetic pathway of PS in mammalian systems. There are two independent pathways that employ different substrates and enzymes for PS synthesis. PSD, PS decarboxylase; PSSI, PS synthase I; PSSII, PS synthase II. **B & C.** Representative immunoblots showing knockdown efficiencies of mouse PSSI (B) and PSSII (C) by shRNA candidates. Myc-tagged PSSI (or PSSII) and the corresponding shRNAs were co-transfected into CHO cells. At 24∼48 hrs after transfection, proteins were extracted and probed by the anti-Myc antibody (∼90 kDa). **D.** Immunoblots of biotinylation experiments detecting the surface (S) and total (T) levels of endogenous ASIC1a in PSSI- and PSSII-knockdown rat cortical neurons. Upper, representative blots. shRNAs targeting PSSI or PSSII were transfected on DIV7. On DIV15, cells were incubated with NHS-biotin (0.25 mg/ml) for 30 min and then lysed. Biotinylated surface proteins were pull-downed with NeutrAvidin agarose beads and immunoblotted, along with the total extracts, for ASIC1a (∼72 kDa) and GAPDH (∼35 kDa). Bottom, quantification of surface/total ratios normalized to the control group. Bars indicate means ± SEM, n = 3. \*\*\**p* < 0.001 (one-way ANOVA with Fisher’s LSD post hoc). **E.** Changes of PS level in PSSI knockdown cells. Staurosporine (1 μg/mL) was applied to shRNA-transfected CHO cells for 2 hrs to induce PS externalization. Externalized PS was then recognized by FITC-Annexin V, and shRNA-transfected cells were visualized by DsRed fluorescence. Left, representative images; right, quantification of FITC intensity in DsRed-positive (transfected) and negative (untransfected) cells. Bars indicate means ± SEM, n = 80 cells for each group. \*\*\**p* < 0.001 (paired *t*-test). **F.** Representative immunoblots for ASIC1a surface (S) and total (T) levels in CHO cells co-transfected with ASIC1a and PSSI shRNAs. EGFP plasmid was co-transfected as a negative control for shRNA targeting (EGFP : ASIC1a : shRNA = 1 : 2 : 3). At 24∼36 hrs after transfection, intact CHO cells were surface biotinylated and processed as in (D) and then immunoblotted for EGFP (∼28 kDa), EGFP-ASIC1a (∼95 kDa) and GAPDH (∼35 kDa). **G.** Biotinylation experiments detecting the surface (S) and total (T) levels of endogenous ASIC1a as in (D) in DIV14 rat cortical neurons treated by different concentrations of DHA for 24 hrs. Left, representative blots. Right, quantification of surface/total ratios normalized to the control (0.1 µM DHA) group. Bars indicate means ± SEM, n = 4. \**p* < 0.05, \*\**p* = 0.0016 (one-way ANOVA with Fisher’s LSD test). **H.** Representative immunoblots for surface (S) and total (T) levels of endogenous ASIC1a on DIV15 rat cortical neurons treated with 10 μM DHA for 24 and 48 hrs.

PSSI was the first identified and the relatively more important PS synthase in rodent cells to date (*1, 49*). A loss-of-function mutation of this enzyme caused 70% reduction of PS in CHO-derived PSA-3 cells (*50, 51*). We further transfected shPSSI- #456 into CHO cells and used Annexin V surface labeling assay to validate the change of PS level in an apoptotic cell model (*11, 52*). In staurosporine-treated CHO cells, we observed a significant decrease of Annexin V signals on shPSSI-#456 transfected cells compared to shRNA-control cells, without any change in the percentage of Annexin V-positive cells (**Fig. 3E**), indicating that PS level was significantly depleted by the knockdown of endogenous PSSI. Next, to find how PS depletion affects ASIC1a surface expression, we co-transfected ASIC1a and shPSSIs into CHO cells (**Fig. 3F**). Interestingly, while the total level of heterologous ASIC1a protein was not changed, the surface ASIC1a protein was significantly reduced in the shRNA-transfected group compared to the control group.

To upregulate PS levels, we applied docosahexaenoic acid (DHA), an important biosynthetic substrate of PS in the nervous system, to cultured cortical neurons (*53*). DHA dose-dependently and time-dependently facilitated the surface expression of endogenous ASIC1a, with the most effective concentration being 3-10 μM **(Fig. 3, G and H)**, indicating that PS regulates surface ASIC1a in a bidirectional manner. Taken together, the above results demonstrate that PSSI activity and PS levels strongly impact ASIC1a surface expression in neurons, suggesting that PS helps to either recruit ASIC1a to or enhance its stability on the surface membrane.

### ASIC1a binds to PS through electrostatic interactions

There are two major types of binding modes between phospholipids and membrane proteins: head-group recognition (for example, between PH domain of PLCδ_1_ and PI(4, 5)P_2_ (*54*), FYVE domain and PI(3)P (*55*), and C2 domain of lactadherin/Annexin V and PS (*1*)) and charge-based interaction (such as that between TRP box and PI(4, 5)P_2_ (*56, 57*)). Because there is no obvious similarity between the sequence of ASIC1a C-terminus and well-known binding motifs for PS, we hypothesized that the polycationic region in the C-terminus of ASIC1a may be responsible for the interaction with the anionic phospholipid in a charge-based manner. To test this hypothesis, we performed alanine scanning starting with the GST-Ct_ASIC1a_ fusion protein, followed by the lipid strip-binding assay (**Fig. 4A**). A di-arginine motif (^467^RR^468^) was found to be critical for binding to PS, as the alanine substitutions of these two (or plus the surrounding L^465^, C^466^, and G^469^) residues abolished the binding of GST-Ct_ASIC1a_ to PS. Neutralization of these two residues by glutamine (Q) substitutions, which largely retained the side chain size, also completely abolished the binding to PS/PA, while lysine (K) substitutions allowed binding to various negatively-charged phospholipids, indicating that the positive charges at these two positions are crucial for binding to phospholipids and the guanidinium groups of the di-arginine motif define the selectivity to PA/PS. We further used pull-down assay with PS-conjugated agarose beads to verify the function of the di-arginine motif in the lipid-protein interaction and the results clearly showed that the conjugated PS preferred binding to wild-type GST-Ct_ASIC1a_ rather than GST-Ct_ASIC1a_-^467^QQ^468^, which was pulled down by PS similarly as GST alone (**Fig. 4B**). Together, these results demonstrate that the positively charged guanidinium groups of ^467^RR^468^ in ASIC1a are essential for the specific binding to PS. To further define the functional significance of the net charges in the di-arginine lipid-binding motif of ASIC1a, we mutated the two arginine residues to various amino acids with neutral (glutamine, ^467^QQ^468^), acidic (glutamate, ^467^EE^468^), or another basic side chain (lysine, ^467^KK^468^) in the full-length ASIC1a. In CHO cells that expressed these constructs, surface biotinylation experiments revealed that the basic-to-neutral and basic-to-acidic mutations markedly decreased the surface levels of ASIC1a in a progressive fashion, while the substitution by lysine (^467^KK^468^) had only moderate impact (**Fig. 4C**). Therefore, the electrostatic state of the lipid-binding motif is a strong determinant for the membrane expression of ASIC1a. Consistent with the decreased surface expression, cells expressing ASIC1a-^467^QQ^468^ and ASIC1a-^467^EE^468^ also exhibited markedly reduced proton-evoked current (*I*_pH6.0_) when exposed to an acidic solution of pH 6.0 (**Fig. 4D**) and such a reduction was insignificant in cells that expressed ASIC1a-^467^KK^468^ when compared to the wild-type ASIC1a. On the other hand, because the ^467^EE^468^ and ^467^QQ^468^ mutants still responded to acid stimulation and the changes in *I*_pH6.0_ correlated with the alterations in the surface expression, the open probability of the channel was unlikely affected by these mutations (**Fig. 4D**). Interestingly, however, the activation time and desensitization time constant (τ) of the ^467^EE^468^ mutant were significantly slower than the wild-type channel, indicating that the lipid binding has a modulatory effect on channel opening kinetics (**Fig. 4, E and F**). This phenomenon is reminiscent of what has been found in P2X channels, where phospholipid binding also accelerates the channel kinetics (*58, 59*).

**Figure 4.**
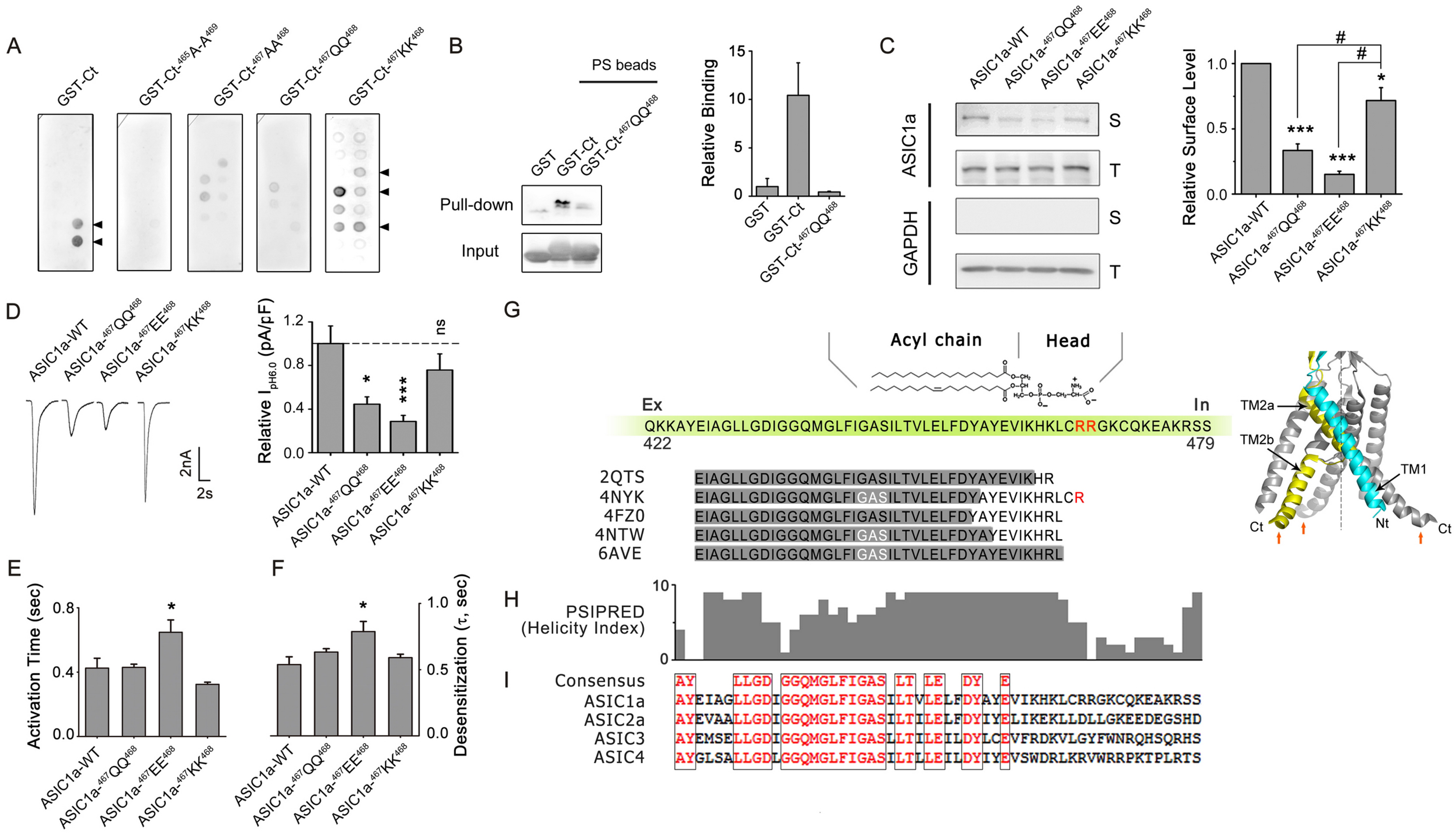
Electrostatic interaction between acidic PS and a di-arginine motif of ASIC1a. **A.** Identification of a di-arginine motif in the proximal region of ASIC1a C-terminus that interacts with PS. Alanine (A), glutamine (Q), but not lysine (K) substitution at this motif abolished binding to anionic phospholipids. **B.** Lipid-protein pull-down assay for GST-Ct_ASIC1a_ containing wild type and the charge-neutralizing mutant sequences of the di-arginine motif, with GST as the negative control (left, representative blots; right, summary). **C.** Biotinylation experiments detecting the surface (S) and total (T) levels of wild type (WT) ASIC1a and its mutants as indicated. CHO cells were transfected with the indicated ASIC1a constructs and biotinylation assay performed at 24∼36 hrs after transfection. Left, representative immunoblots. GAPDH was used as a cytoplasmic protein control. EGFP-ASIC1a, ∼95 kDa; GAPDH, ∼35 kDa. Right, quantification of surface/total ratios ASIC1a normalized to the WT group. Bars indicate means ± SEM, n = 4. \**p* < 0.05, \*\*\**p* < 0.001 (vs. Ctrl); ^#^*p* < 0.05 (vs. ^467^KK^468^ group) (one-way ANOVA with Fisher’s LSD test). **D.** Electrophysiological functions of WT ASIC1a and its mutants of the di-arginine motif. Whole-cell recordings were performed on CHO cells transfected with the indicated ASIC1a constructs 1 day after transfection. Cells were held at −60 mV while a pH 6.0 solution was applied via perfusion. Left, representative current traces; right, summary of peak current density for n = 10 cells of WT, 7 cells of ^467^QQ^468^, 10 cells of ^467^EE^468^, and 6 cells of ^467^KK^468^. Bars indicate means ± SEM. \**p* = 0.0154, \*\*\**p* < 0.001 (one-way ANOVA with Fisher’s LSD test). **E & F.** Quantifications of the activation rate (E) and desensitization rate (F) of acid (pH 6.0)-evoked currents recorded in (D). \**p* < 0.05 (one-way ANOVA with Tukey’s post hoc). **G.** Schematic diagram of interaction between PS negative head group and ^467^RR^468^ motif of ASIC1a at membrane-cytosol interface. The sequence marked in green encompasses transmembrane domain II (TMII) and beginning of the cytoplasmic C-terminus of ASIC1a, with the di-arginine motif indicated in red. TMII sequences revealed in the resolved chicken ASIC1 crystal structures are shown in the alignment below. Right, ribbon view of ASIC1a structure with transmembrane domains and short intracellular stretches. TMI and TMII connecting to N- and C-termini of one subunit are shown in blue and yellow, respectively. The structural model of ASIC1a from PDB 6AVE was represented by PyMOL v0.99rc6. The positions of the di-arginine motifs are indicated by red arrows. **H.** Helicity index for TMII and C-terminus of ASIC1a (0–10, generated by PSIPRED). Residues are aligned with (G). **I.** Alignment of TMII helices and proximal C-terminal domains among human ASIC family members.

Interestingly, although the majority of published ASIC1 structures do not show cytoplasmic N- and C-termini, the following analyses suggest that the di-arginine motif is strategically situated near the inner leaflet of PM, where it can easily interact with the negatively charged phospholipid head group. First, both the resolved ASIC1 structures (**Fig. 4G and fig. S7A**) and hydrophobicity analyses using several server-based programs and algorithms (*60*) **(fig. S7B**) indicated that the two arginines are within the initial segment of the cytoplasmic C-terminus immediately downstream from the second transmembrane (TMII) domain. Second, the crystal structure of chicken ASIC1 (*61*) shows the existence of a hinge at the GAS motif (important for ion selectivity of ASIC1a) within TMII, which makes the proximal C-terminus more parallel to the membrane, placing the arginines even closer to membrane lipids in the inner leaflet (**Fig. 4G, right, and fig. S7A, right**). Third, helicity analysis using PSIPRED 3.3 showed that the helix moment of TMII segment (between I^428^ and L^465^) terminates abruptly at ^467^RR^468^, suggesting the lipid binding motif to be likely in a disordered region (**Fig. 4H**). Fourth, amino acid sequence alignments of related ASICs and ENaC isoforms, as well as ASIC1a from different species, clearly indicate that the high degree of conservation of TMII ends right at the beginning of the C-termini, where the di-arginine motif is only present in ASIC1 isoforms (**Fig. 4I and fig. S7C**). Altogether, these analyses suggest that the di-arginine motif is located in a flexible region immediately adjacent to the membrane-cytosol interface, with easy access to anionic phospholipids in PM inner leaflet to form electrostatic interactions, which appear to support the cell surface expression of ASIC1a.

### ASIC1a preferentially traffics with PS in endocytic organelles to the leading edge

PS has been shown to be associated with Cdc42 and contribute to the formation of cell polarity in yeast (*5*). In CHO cells, we also found cytosolic Lact-C2 to have a polarized distribution pattern. Although CHO cells do not show complex protrusions, the overexpressed Lact-C2 typically accumulated in a small number of peripheral areas of these cells. By incubating the CHO cells with FITC conjugate of cholera toxin B subunit (FITC-CTxB), the lipid raft marker that labels membrane microdomains enriched with cholesterol and GM1 ganglioside receptors, we interestingly found that at least some of the Lact-C2 hotspots strongly overlapped with lipid rafts at the cell edge (*9, 62*) (**Fig. 5A, left**). By co-expressing Lact-C2 with Rho family members, Cdc42 and Rac1, which are usually concentrated at filopodia and lamellipodia of fibroblasts (*63*), we found colocalization of Lact-C2 with Cdc42 and Rac1 at the leading edge of CHO cells **(Fig. 5A, middle and right)**. When co-expressed with ASIC1a, Lact-C2 was found to be strongly colocalized with ASIC1a not only at the edge of cell membrane, but also in vesicular structures inside the cell **(Fig. 5B)**. As a negative control, the Lact-C2 mutant unable to bind to PS, Lact-C2-AAA, was more evenly distributed throughout the cytoplasm and nucleus of the cell and to a large extent did not colocalize with ASIC1a **(Fig. 5, B and C)**.

**Figure 5.**
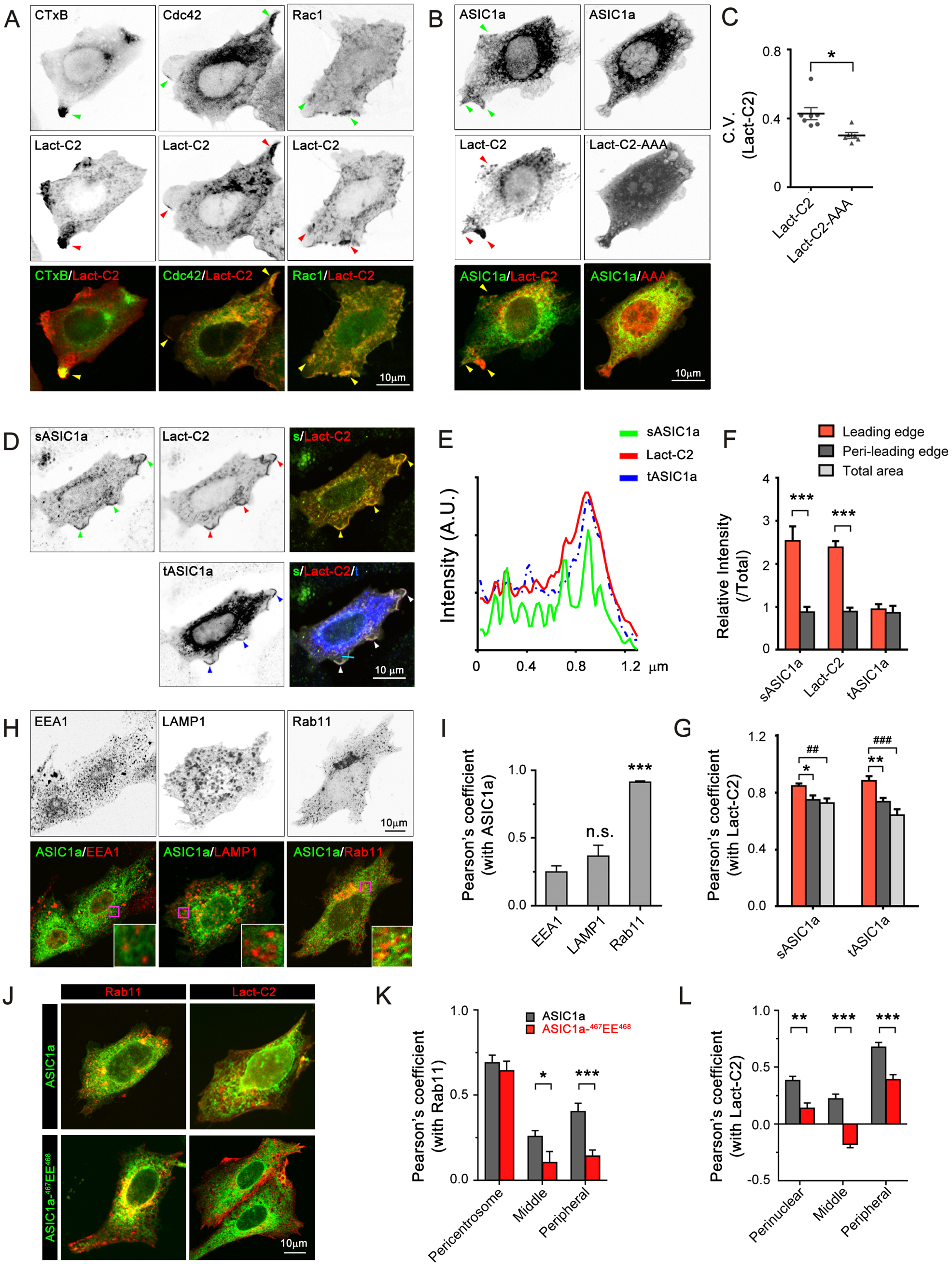
PS-binding is required for normal membrane targeting of ASIC1a to cell leading edge. **A.** Representative confocal images of CHO cells transfected with mCherry-Lact-C2 and EGFP-Cdc42 (middle) or EGFP-Rac1 (right). After fixation, cells were labeled with FITC-CTxB (left), or immunostained with antibodies of the corresponding fluorescent proteins (middle and right). Arrowheads, leading edges or hotspots enriched of Lact-C2 and Rho family members. **B.** Representative confocal images of CHO cells transfected with EGFP-ASIC1a and mCherry-Lact-C2 (left) or the PS-binding deficient mutant, mCherry-Lact-C2-AAA (right). Arrowheads, leading edges or intercellular vesicles enriched of Lact-C2 and ASIC1a. **C.** Comparison of distributions between wild type Lact-C2 and Lact-C2-AAA in CHO cells. Signal variability of each group is indicated by coefficient of variation (C.V.). **D.** Representative confocal images of CHO cells transfected with N-terminal EGFP-tagged HA-ASIC1a and mCherry-Lact-C2. After fixation, cells were immunostained by anti-HA (for surface ASIC1a, sASIC1a or s) before permeabilization, and then anti-EGFP (for total ASIC1a, tASIC1a or t), and anti-mCherry (for Lact-C2) after permeabilization. Arrowheads, leading edges. Blue line indicates line scan area in (E). **E.** Intensity profiling of line scan through a leading edge in (D, lower right panel, blue line). **F.** Quantification of distributions of indicated proteins in different compartments of CHO cells in (D). Fluorescent intensity of the indicated protein in each area was normalized to the mean intensity of the same protein in the entire cell area. n = 11 cells, \*\*\**p* < 0.001 (Student’s *t*-test). **G.** Pearson’s correlation coefficients between Lact-C2 and surface ASIC1a or total ASIC1a in different areas of CHO cells in (D). n = 11 cells. \**p* < 0.05, \*\**p* < 0.01, ^##^ *p* < 0.01, ^###^*p* < 0.001 (one-way ANOVA with Fisher’s LSD test). **H.** Screening for endosomal pool of ASIC1a. CHO cells were co-transfected with EGFP-ASIC1a and mCherry-Lamp1 (middle) or mCherry-Rab11a (right) and fixed at 24 hrs after transfection. Anti-EEA1 antibody was used to label early endosomes (left). Insets show the enlarged views of the pink boxes. **I.** Pearson’s correlation coefficients between ASIC1a and endosomal markers as in (H). n = 5 cells for each group. n.s., no significance; \*\*\**p* < 0.001, vs. EEA1 group (one-way ANOWA with Fisher’s LSD post hoc). **J.** Representative confocal images of CHO cells transfected with wild type EGFP-ASIC1a (top) or its ^467^EE^468^ mutant (bottom) and endosomal marker mCherry-Rab11a (left) or mCherry-Lact-C2 (right). After fixation, cells were immunostained with corresponding antibodies for each fluorescent protein. Note, despite the high enrichment in recycling endosomes for both markers, Rab11a is more concentrated at pericentrosomal zone than Lact-C2, which is more preferentially distributed at peripheral of the cells. **K & L.** Pearson’s correlation coefficients between Rab11a (K) or Lact-C2 (L) and ASIC1a or its ^467^EE^468^ mutant as in (J). n = 8 cells for each group. \**p* = 0.0248, \*\**p* < 0.01, \*\*\**p* < 0.001 (Student’s *t*-test). Middle, the area in the middle between the cell edge and centrosome (K) or nuclear (L).

Not surprisingly, when we used HA-ASIC1a to detect the surface fraction of ASIC1a by staining non-permeabilized cells with the anti-HA antibody, we found that the surface ASIC1a was also highly concentrated at the leading edge of CHO cells and strongly overlapped with Lact-C2 (**Fig. 5, D to G, and fig. S8**). Line scan through the leading edge of CHO cells on the Z-max-projection images showed that the peaks of surface ASIC1a colocalized strongly with that of Lact-C2, even much better than that of total ASIC1a (**Fig. 5E**). These results further demonstrate that surface ASIC1a is bound to PS to acquire a strongly polarized cellular distribution and function at the highly dynamic front edge.

To study the relationship between ASIC1a and PS during vesicular transport, we used multiple endosomal markers to identify the intracellular pool(s) of ASIC1a (*26, 64*). We found ASIC1a to be more preferentially colocalized with Rab11a than EEA1 (early endosome) or Lamp1 (lysosome), indicating that a large fraction of the intracellular ASIC1a exists in recycling endosomes (RE) (**Fig. 5, H and I**). This result is particularly interesting because RE is also the largest endosomal pool of PS (*8, 40, 65*) where the lipid contributes to vesicular trafficking (*66, 67*).

Next, to learn the role of PS binding in intracellular trafficking of ASIC1a, we compared the distribution patterns of wild-type ASIC1a and ASIC1a-^467^EE^468^. We separated the cellular regions into perinuclear, middle, and peripheral areas. Although both ASIC1a-^467^EE^468^ and wild-type ASIC1a were abundantly present in pericentrosomal Rab11a-positive (Rab11a^+^) vesicles, the amounts of ASIC1a-^467^EE^468^ associated with the middle and peripheral Rab11a^+^ vesicles were markedly decreased as compared to wild-type ASIC1a (**Fig. 5, J and K**). Furthermore, Lact-C2 displayed more pronounced peripheral distribution than Rab11a, where it showed excellent colocalization with wild-type ASIC1a, which was dramatically reduced with ASIC1a- ^467^EE^468^ (**Fig. 5, J and L**). Because PS is mainly enriched in the PM and endocytic membranes, and its concentration is gradually increased from RE to the inner leaflet of PM (*68*), our results suggest that uncoupling with PS also impairs endocytic trafficking of ASIC1a-^467^EE^468^ through the PS-dependent pathway. Together, the above data suggest that RE is the largest endosomal pool for both ASIC1a and PS, and normal vesicular transport of ASIC1a relies on binding to PS (*69*).

### Uncoupling ASIC1a-PS binding causes mistargeting and dysfunction of ASIC1a

In neuronal cultures, we used two methods to determine the physiological relevance of PS-binding on ASIC1a. First, we expressed the PS-binding deficient mutant, ASIC1a- ^467^EE^468^, into *Asic1a*^-/-^ mouse neurons (for a clean genetic background) and evaluated the changes in ASIC1a channel function. In *Asic1a*^-/-^ cortical neurons untransfected or transfected with the control cDNA for GFP, patch clamp recordings showed almost no detectable *I*_pH6.0_, which was successfully restored by the expression of wild-type ASIC1a (**Fig. 6A**). The expression of ASIC1a-^467^EE^468^ also restored *I*_pH6.0_, but the current density was much smaller than the expression of wild-type ASIC1a. On the other hand, ASIC1a-^467^KK^468^ restored *I*_pH6.0_ to a similar degree as the wild-type channel (**Fig. 6A**). Furthermore, when expressed in *Asic1a*^-/-^ mouse neurons, the activation time and desensitization rate of the ^467^EE^468^ mutant were significantly prolonged compared to the wild-type channel (**Fig. 6, B and C**). These findings are entirely consistent with the results obtained from CHO cells (**Fig. 4, D to F**), further supporting the idea that phospholipid binding not only facilitates surface expression of ASIC1a but also has a regulatory effect on the channel opening kinetics in neurons.

**Figure 6.**
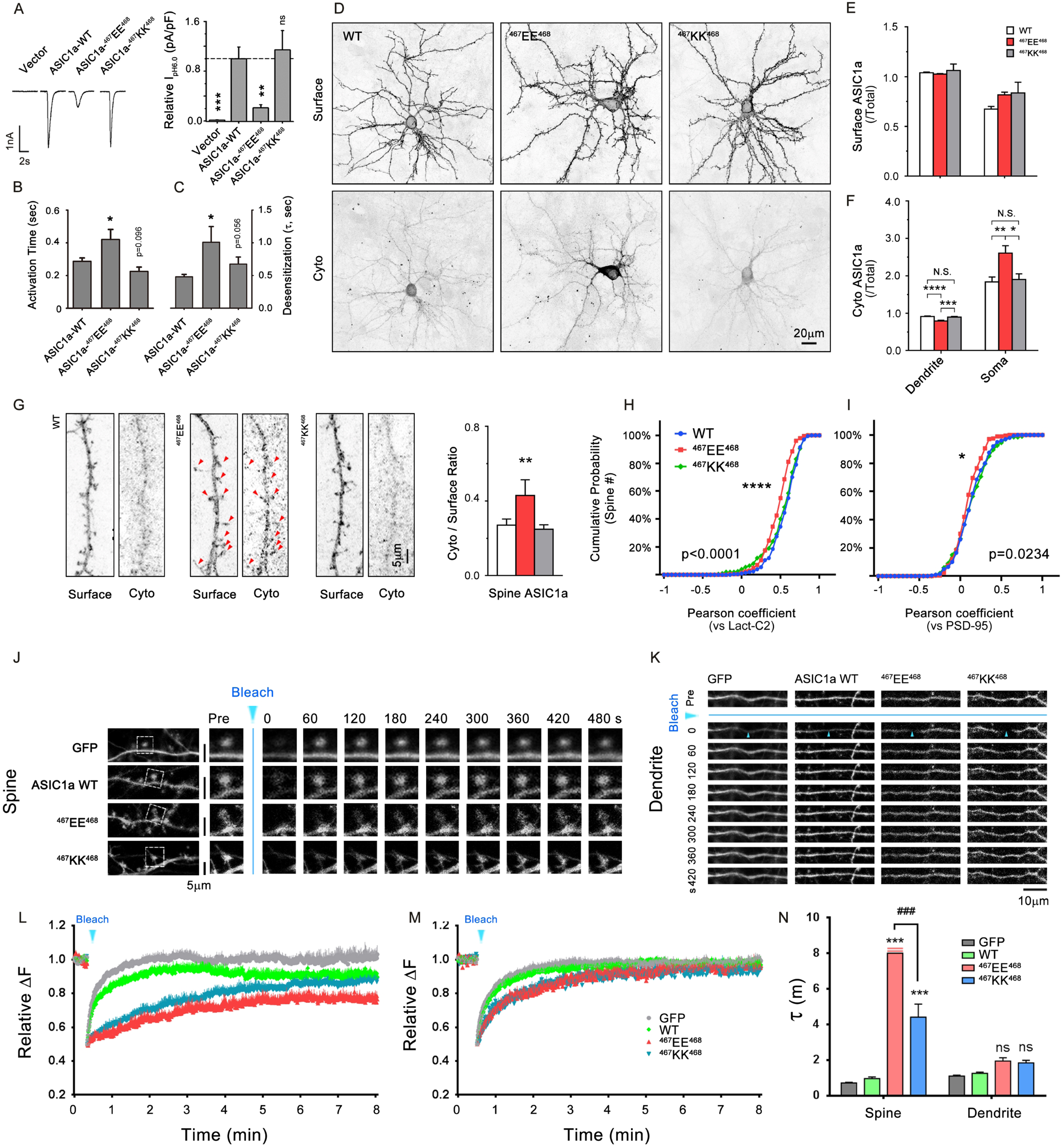
PS-binding controls synaptic targeting of ASIC1a. **A**. Acid-evoked currents in mouse *Asic1a*^-/-^ cortical neurons transfected with GFP control, wild type (WT) ASIC1a or its di-arginine motif mutants at DIV7-10. Whole-cell voltage clamp recordings were performed at least 48 hrs after transfection, with the cells held at −60 mV and current evoked by perfusion with a pH 6.0 solution. Left, representative current traces; right, summary of peak current density. Bars indicate means ± SEM, n = 10 for each group. \*\**p* = 0.0038, \*\*\**p* = 0.0004 vs. WT ASIC1a (one-way ANOVA with Fisher’s LSD post hoc). **B & C.** Quantification of activation rate (B) and desensitization rate (C) of currents recorded in (A). \**p* = 0.0134, \*\**p* = 0.0070, vs. WT ASIC1a (one-way ANOVA with Fisher’s LSD post hoc). **D.** Representative confocal images of DIV28 rat cortical neurons transfected with WT HA-ASIC1a, HA-ASIC1a-^467^EE^468^, or HA-ASIC1a-^467^KK^468^ mutant. After fixation, surface ASIC1a was immunostained with the monoclonal HA antibody under non-permeabilized conditions and visualized by Alexa Fluor 488-conjugated secondary antibody (upper), and cytoplasmic (Cyto) ASIC1a was detected with the polyclonal HA antibody and Alexa Fluor 568-conjugated secondary antibody after permeabilization (lower). **E & F.** Quantification of mean intensity ratios of dendritic or somatic ASIC1a to total ASIC1a in surface (E) and cytoplasm (F) of DIV28 rat cortical neurons in (D). n = 10 cells for each. For cytoplasmic ASIC1a (bottom panel), n.s., no significance, \**p* < 0.05, \*\**p* < 0.01, \*\*\**p* < 0.001, \*\*\*\**p* < 0.0001 (one-way ANOVA with Tukey’s post hoc). **G.** Representative confocal images of neuronal dendrites stained the same way as in (D) and quantification of cytoplasmic/surface ratios of ASIC1a intensity at the spine head areas. n = 100 spines from 10 cells each. \*\**p* < 0.01, vs. WT, one-way ANOVA with Turkey’s post hoc; WT vs. ^467^KK^468^, no significance. **H-I.** Cumulative probabilities of Pearson’s correlation coefficients between surface ASIC1a and Lact-C2 (H), or PSD-95 (I) on the SIM z stack maximum projection images of dendritic spines, n = 200 spines from 10 cells for each group. H. \*\*\*\**p* < 0.0001, one-way ANOVA with Turkey’s post hoc; WT vs. ^467^EE^468^, \*\*\*\**p* < 0.0001; ^467^EE^468^ vs. ^467^KK^468^, \*\**p* < 0.01; WT vs. ^467^KK^468^, no significance. I. \**p* = 0.0234, one-way ANOVA with Turkey’s post hoc; WT vs. ^467^EE^468^, no significance; ^467^EE^468^ vs. ^467^KK^468^, \**p* < 0.05; WT vs. ^467^KK^468^, no significance. **J.** Fluorescence recovery after photobleaching (FRAP) analysis on DIV21 for cortical neurons transfected with EGFP control or EGFP-ASIC1a of WT or mutants. Photobleaching was performed in a small region (less than 1 μm^2^) located at the spine head by a 488-nm laser and time-lapse imaging was acquired before and after bleaching. Left, representative images of example dendrites; Blue arrowhead, photobleaching for 10 s. Right, high-magnification images of the photobleached spines (white squares in the left panel) acquired at a fixed interval (60 s) after bleaching. **K.** FRAP analysis on the dendritic shaft areas performed similarly as in (J). Small up-pointing blue arrow-heads, photobleached dendritic areas. **L & M.** Time courses of fluorescence recovery on the spine head and dendritic shaft areas recorded as in (J) and (K); baseline before photobleaching, 20 s. L, n = 6, 7, 7, and 4, M, n = 6, 6, 8, and 8, for GFP, ASIC1a WT, ^465^EE^467^, and ^465^KK^467^ groups, respectively. **N.** Quantification of time constants (τ) of FRAP shown in (L) and (M). \*\*\**p* < 0.001; ns, no significance, vs. ASIC1a WT; ^###^*p* < 0.001, vs. ^467^EE^468^ (one-way ANOVA with Turkey’s LSD post hoc). For the ^467^EE^468^ group in spines, τ was determined to be > 8 min, but τ = 8 min was used for calculation.

Next, we examined how the charge-reversal mutation of the lipid binding motif impacted surface ASIC1a by immunocytochemistry **(Fig. 6D)**. In rat cortical neurons that expressed various ASIC1a constructs with extracellular HA-tag, the surface fraction of HA-ASIC1a was bound by the monoclonal anti-HA antibody under non-permeabilized conditions, which was followed by permeabilization and detecting the cytoplasmic fraction of HA-ASIC1a with the polyclonal anti-HA antibody. In order to ensure descent fluorescent signals for quantification, we only analyzed cells that had strong surface staining signals. Despite equivalent surface levels for neurons that expressed HA-tagged wild-type ASIC1a, ASIC1a-^467^EE^468^, and ASIC1a-^467^KK^468^, the level for the cytoplasmic fraction of the ^467^EE^468^ mutant in the soma was significantly higher than that of the wild-type and the ^467^KK^468^ mutant (**Fig. 6, D to F**), consistent with the biotinylation results showing decreased surface expression of ASIC1a-^467^EE^468^ (**Fig. 4C**). Notably, a closer examination of the spine heads revealed that ASIC1a- ^467^EE^468^ accumulated in the cytoplasm just underneath the surface of spine heads; this was in stark contrast with the wild-type ASIC1a, which showed less than 6% retention in the cytoplasmic space of the spine (**Fig. 6, F and G, Fig. 2L, and fig. S6C**). More interestingly, even for the surface ASIC1a-^467^EE^468^, the colocalization with Lact-C2, or PSD-95, was also poorer than the surface wild-type ASIC1a and the ^467^KK^468^ mutant, implicating mistargeting of ASIC1a-^467^EE^468^ in synapses (**Fig. 6, H and I, and fig. S9**). To further test the function of PS-binding motif on synaptic targeting of ASIC1a proteins in live neurons, we also employed time-lapse imaging and fluorescent recovery after bleaching (FRAP) techniques on EGFP-ASIC1a-transfected cortical neurons (*18, 28*). The recovery of photobleached fluorescent signals presents the diffusion or transport processes of the fusion proteins in a specific cellular compartment. At the spine head areas, the recovery time constant (τ) of wild-type ASIC1a protein is 0.96 ± 0.09 min, a little slower than the GFP control (0.72 ± 0.04 min). In contrast to wild-type ASIC1a, ASIC1a-^467^EE^468^ mutant showed significantly delayed recovery rate (τ > 8 min), while ASIC1a-^467^KK^468^ mutant partially rescued the time constant to 4.41 ± 0.73 min (**Fig. 6, J and L**), indicating that the positively charged guanidinium groups in the PS-binding motif are critical for the synaptic targeting efficiency of ASIC1a protein. On the other hand, the transporting efficiencies of the ASIC1a mutants in dendritic shaft, where protein transport process is more dominant than protein sorting and targeting, were comparable to that of the wild-type ASIC1a (recovery time constants: GFP, 1.11 ± 0.05 min; wild type ASIC1a, 1.26 ± 0.07 min; ^465^EE^467^, 1.96 ± 0.18 min; ^465^KK^467^ 1.84 ± 0.15 min, *P* > 0.5 *vs*. wild-type ASIC1a for all) (**Fig. 6, K, M, N**). Therefore, the PS-binding does not influence the dendritic transport of ASIC1a. Together, these results show that PS is more important for the spine distribution and synaptic delivery of ASIC1a than dendritic transport of the channel.

We then used an interfering peptide that mimics the ASIC1a PS-binding motif to uncouple the endogenous ASIC1a and PS. We designed a short peptide that contains a cell membrane-penetrating sequence TAT (YGRKKRRQRRR), a C-terminal fragment of ASIC1a flanking the ^467^RR^468^ motif (LCRRGKCQKE) and a CAAX-box prenylation sequence of K-Ras (KSKTKCVIM) **(Fig. 7A)** (*70, 71*). Application of this peptide to cultured cortical neurons led to a significant reduction of surface ASIC1a, without altering the total protein level, indicating that the peptide successfully competed PS off the endogenous ASIC1a (**Fig. 7B**). The ability of the peptide to reduce surface ASIC1a was lost by substituting the arginines (RR) with glutamates (EE) or glutamines (QQ), but partially retained by the substitution with lysines (KK) (**Fig. 7B**), further demonstrating that the positive charges in ASIC1a C-terminal motif were essential for PS binding. We also tested a series of peptides without the CAAX surface targeting motif and found them to have no effect on surface expression of the endogenous ASIC1a (data not shown), indicating that lipid binding by the basic motif only occurs near the membrane inner leaflet.

**Figure 7.**
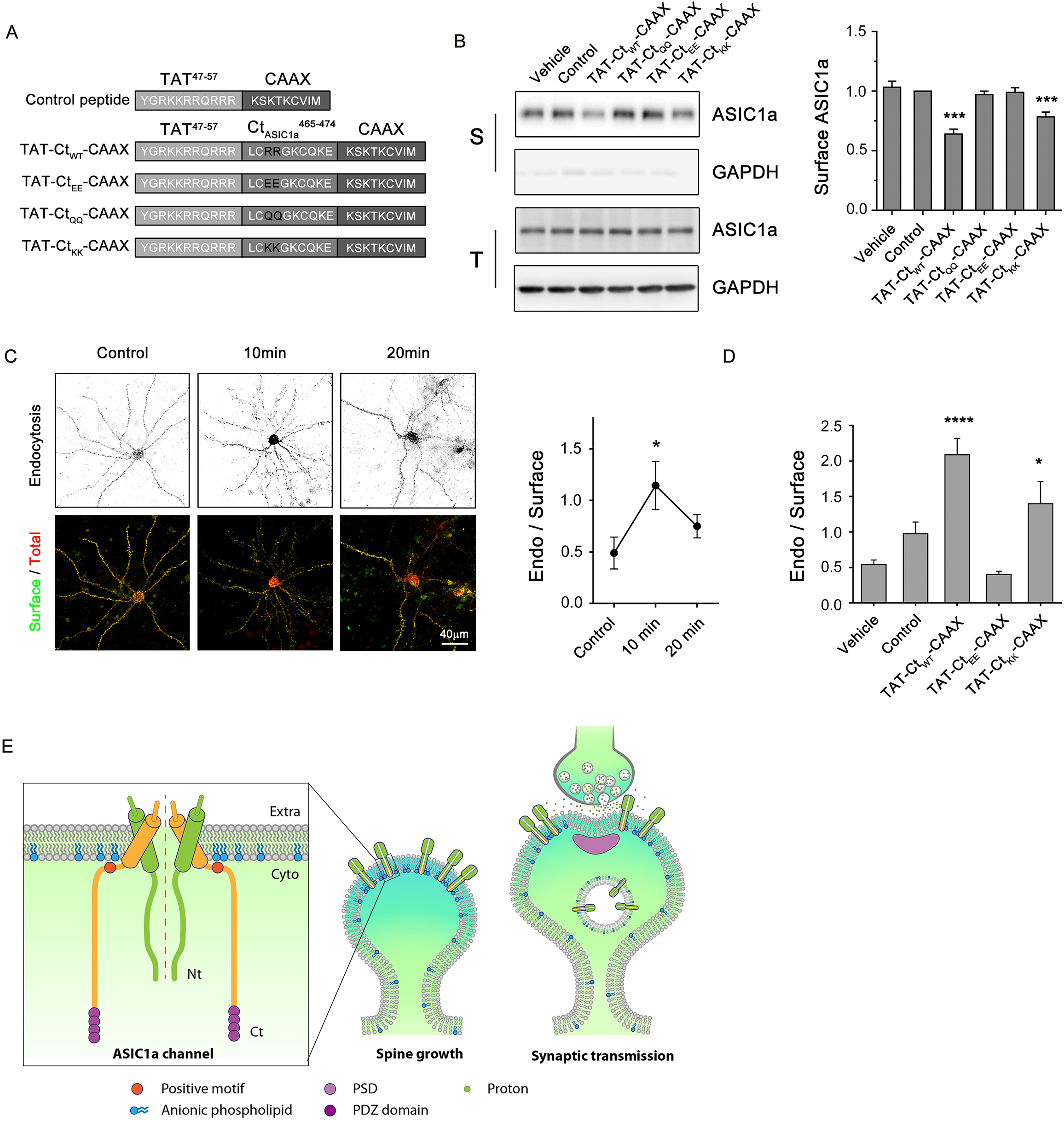
Disrupting ASIC1a binding to PS enhances channel internalization. **A.** Schematic diagram of cell-penetrating peptides designed to compete with the di-arginine motif of ASIC1a for PS binding. **B.** Surface biotinylation analysis detecting the surface (S) and total (T) levels of endogenous ASIC1a in rat cortical neurons treated with 5 μM of the cell-penetrating peptides shown in (A) on DIV14-21. After 24 hrs, cells were subject to surface biotinylation, and proteins were extracted, precipitated with the NeutrAvidin agarose beads, and immunoblotted for ASIC1a (∼72 kDa) and GAPDH (∼35 kDa). Left, representative blots; right, quantification of the surface/total intensity ratio of each band normalized to the control. Bars indicate means ± SEM, n = 5-10 for each group. \*\*\**p* < 0.001, vs. control group (one-way ANOVA with Fisher’s LSD post hoc). **C & D.** Antibody feeding assay showing ASIC1a endocytosis in response to peptide application. Surface HA-ASIC1a protein expressed in rat cortical neurons transfected with N-terminal EGFP-tagged HA-ASIC1a was labeled alive by the HA polycloncal antibody for 10 min before the TAT-Ct_WT_-CAAX (C) and other targeting peptides (D, all at 10 μM) were added for 10 and 20 min (C) or 10 min (D). Internalized and total HA-ASIC1a were probed after cell fixation and permeabilization as described in Methods. n = 6-11 cells for each group. \**p* < 0.05, \*\*\*\**p* < 0.0001, vs. vehicle group (one-way ANOVA with Turkey’s post hoc). **E.** Schematic diagram of ASIC1a-PS interaction at the synaptic membrane of the dendritic spine. ASIC1a and PS are highly enriched in dendritic spine head area and recycling endosomes. ASIC1a interacts with anionic phospholipids, such as PS, via a positively charged di-arginine motif. The perisynaptic localization of ASIC1a in mature synapses is illustrated at right, which supports the idea of ASIC1a activation by the spill-over of protons co-released from synaptic vesicles with neurotransmitters during high frequency and/or strong neuronal activities. Extra: extracellular; Cyto: cytoplasmic.

By using the competing peptide, we asked whether in addition to facilitating ASIC1a recycling, PS binding also stabilizes surface-expressed ASIC1a. In an anti-HA antibody feeding assay, application of 10 μM wild-type (RR) peptide dramatically increased the amount of surface-bound anti-HA antibody in endocytic vesicles over time (**Fig. 7, C and D**). The endocytosis/surface ratio increased from the basal level of 0.49 ± 0.15 to 1.14 ± 0.23 after 10 min of the peptide treatment and then declined slightly at 20 min (**Fig, 7C**), indicating a transient acceleration of surface ASIC1a internalization. By contrast, the ^467^EE^468^ peptide failed to cause HA-ASIC1a endocytosis and, if anything, it might have reduced the internalization as compared to the vehicle or control peptide (**Fig. 7D**). On the other hand, the peptide with positively charged ^467^KK^468^ partially recapitulated the effect of wild-type peptide in accelerating HA-ASIC1a internalization (**Fig. 7D**). These results indicate that PS binding greatly enhances the membrane stability of surface ASIC1a such that when disrupted by a competing peptide, the channel undergoes rapid internalization through endocytosis.

## Discussion

### Asymmetric distribution of PS in neurons

Asymmetric distribution and polarity are the most important characteristics for the functional efficiency of membranous structures in eukaryotic cells, and phospholipids play a central role in their establishment. Previous studies showed that PIP_3_ contributes to cell polarity, motility and chemotaxis (*72*), but the function and importance of other phospholipids in polarity still remain mysterious. The report on asymmetric distribution of PS and its accumulation in protrusions in yeast (*S. cerevisiae*) in the absence of externalization (*5*) prompted our interest in investigating the distribution and function of PS in more complex eukaryotic cells (*73, 74*), especially neurons because of their elaborated structural features such as dendrites and axons, as well as spines and synapses, of which morphological and functional strength are highly correlated to neuronal activities (*75*).

The present study provides the first evidence that the anionic PS is concentrated at the spine head, especially on peri-synaptic sites, which differs from pre- or postsynaptic distributions of other negatively charged phospholipids, such as PIP_2_ (*76*), PIP_3_ (*3, 4*) and PA (*77*), implicating that PS may play a distinct role in recruiting a unique group of synaptic proteins. While the major components of cell membrane, such as PC (∼40%) and PE (20-30%) are all electrically neutral, PS is the most abundant charged phospholipid on the PM. Accumulating evidence shows that negatively charged phospholipids are concentrated at the synapse to help recruit synaptic proteins with positively charged motifs (78, 79). The relative abundance (more than 10% of total membrane phospholipids) of PS and its asymmetric localization to the inner leaflet of PM (reaching ∼30% (*1*)) provide considerable negative charges (*68*), making PS the major contributor of synaptic electronegativity. Our findings support the hypothesis that the postsynaptic membrane contains PS-enriched negatively charged micro- or nano-domains with distinct lipid and protein compositions. Such an organization may be crucial for specific signaling pathways. PS has been previously reported to bind to Cdc42 (*5, 80*), syntaxin-1a (*4*), protein kinase C (PKC) (*81*), c-Raf (*82*) and K-Ras (*83, 84*). In more recent studies, PS was also reported to stabilize and regulate the function of cystic fibrosis transmembrane conductance regulator (CFTR) (*85*) and the Piezo1 channel (*6*).

### Specialized functions of PS and ASIC1a in neurons

Although some studies have reported that lysophosphatidylcholine (LPC) and arachidonic acid (AA) can activate ASIC3 or potentiate acid-induced currents (*86, 87*), phospholipid regulation of ASIC1a has not been shown before. Here, we identified that anionic phospholipids, especially PS, bind and stabilize ASIC1a on the membrane of spine heads of mammalian neurons, where ASIC1a is known to participate in synaptic plasticity. Notably, there is strong evidence that both ASIC1a and PS are highly expressed and specialized in neurons and contribute to normal brain functions: ASIC1a regulates spine components (*88*) and remodeling (*19, 20*), synaptic transmission (*21, 22*), LTP (*23–25, 89*), and fear-related memory (*90*); PS and DHA have long been known to be crucial for human brain development and their deregulation contributes to neurodegeneration and psychiatric diseases (*91, 92*). Recent studies revealed that the gain-of-function mutations of PSSI underlie the human developmental disorder, Lenz-Majewski syndrome (*11*), while the loss-of-function mutations of PSSI may contribute to Nablus mask-like facial syndrome (*10*). Moreover, double knockout of PSSI and PSSII genes causes lethality in mice (*93*), strongly indicating that PS is essential for neural development and normal body function *in vivo*. Results from the present study further reveal this lipid-protein interaction in highly specialized micro-environment, reinforcing the possibility that PS and ASIC1a function together in neurons to regulate synaptic plasticity.

### Spine enrichment and peri-synaptic localization of ASIC1a and PS

Our super-resolution imaging results obtained from two distinct techniques clearly show that both ASIC1a and Lact-C2 are enriched in dendritic spine heads, with a prominent peri-synaptic localization adjacent to the postsynaptic sites where PSD-95 and GluN1 receptor accumulate. This localization pattern is likely true as we took the precaution to avoid artifacts by 1) making certain that heterogeneous expression did not cause saturation such that the postsynaptic density could be filled by the overflow of the exogenous ASIC1a or Lact-C2 proteins, and 2) comparing the distributions of GFP, which diffused normally into the postsynaptic density, and Lact-C2, which did not, indicating that Lact-C2 prefers to stay away from the postsynaptic density. Furthermore, similar distribution patterns have been well-described for other membrane receptors, such as metabotropic glutamate receptors (*94*).

Our findings are consistent with previous reports about spine enrichment of ASIC1a (*19*), but the relationship between ASIC1a and PSD-95 turns out to be different from that suggested by others. It was reported that ASIC2a binds to a PDZ domain of PSD-95, which helps ASIC1a targeting to the dendritic spine by oligomerization (*20*), and a more recent study showed that synaptic ASIC1a activity is inhibited by PSD-95 through ASIC2a integration (*95*). Since *in vivo* analysis demonstrated that ASIC1a expression peaks much earlier than ASIC2a during neurodevelopment, especially during the prenatal stage (*41, 96*), it is more likely that the gradually increased ASIC2a serves to stabilize these channels at the postsynaptic density only at the later developmental stage (such as during the adult period). Combining the present results with the earlier report, we propose that the majority of spine-enriched ASIC1a channels function at the peri-synaptic sites, at least under steady state. This hypothesis does not exclude some specific conditions, such as during neuronal activities, under which the localization of ASIC1a or the relationships between ASIC1a and PSD-95 may change. Furthermore, this peri-synaptic localization suggests that during neurotransmission, ASIC1a is mainly activated by the spill-over of protons co-released with the neurotransmitters. With few exceptions (*97*), ASIC1a has been shown to contribute to LTP induced by high frequency stimulation (HFS) (*25*), suggesting that overflow of protons from the limiting space of synaptic cleft (*98*), which occurs more likely with HFS, is critical for ASIC1a regulation of synaptic strength.

### Electrostatic recognition between PS and ASIC1a

Protein partners interact with PS mainly in two ways: specialized PS-binding motif, such as C2 domain of PKC, Annexin V and Lactadherin, and less selective polycationic stretches, like Ras and Rho family members (*68*), as well as ASIC1a shown in this study. Polybasic domains that bind anionic phospholipids have also been discovered in many other neuronal receptors, including TRP channels (TRP box) (*56, 57*), P2X receptors (*58*) and ENaC (*99, 100*). We found a di-arginine motif at the proximal C-terminus of ASIC1a to be involved in electrostatic interaction with PS. Interestingly, the membrane-proximal stretch of ASIC1a C-terminus, which contains the lipid-binding motif, is also involved in the interaction with AP2 complex (*27*), suggesting that AP2 may facilitate ASIC1a endocytosis by competing with anionic phospholipids for interaction with ASIC1a C-terminus. Furthermore, previous functional studies on chicken ASIC1 (cASIC1, the ortholog of rodent and human ASIC1a) showed that adding Arg466 back to the C-terminal truncation of ASIC1a (ΔASIC1) successfully rescued its structural and functional deficiency (*101, 102*), demonstrating that the positively-charged residue is essential for its membrane stability and normal functions.

### Co-transport in endocytic organelles

Different organelles contain distinct lipid compositions which may contribute to vesicular characteristics and various cellular functions, including size, formation, fusion, protein sorting, and transport processes of intracellular vesicles (*55, 103*). With small negative head groups, anionic PS and PA have been reported to affect vesicle curvature (*104, 105*) and membrane trafficking. In addition to the abundant presence in the inner leaflet of the PM, PS has also been found in cytoplasmic compartments, mainly endocytic organelles (*40, 65*), and to be associated with recycling pathway (*5, 69, 106*). Interestingly, our data demonstrate for the first time that Rab11-positive recycling endosomes are also a large intracellular pool of ASIC1a. Interestingly, although the ASIC1a mutant unable to bind to PS still found its way to accumulate in Rab11-positive RE structures, or even in the spine head, its presence in Lact-C2-positive vesicles is significantly reduced. Because PS is synthesized in luminal monolayer of endoplasmic reticulum, exposed at Middle-to-Late Golgi and enriched in RE and PM (*65*), our data suggest that the trafficking deficit and retention of the ASIC1a-^467^EE^468^ mutant may occur during the transport from RE to PM, where an increasing gradient of PS exists.

In summary, we show that PS is crucial for synaptic localization and function of ASIC1a. ASIC1a binds to PS in the inner leaflet of the PM through a di-arginine motif at its proximal C-terminus, which supports ASIC1a spine targeting and membrane stability (**Fig. 7E**). The spine localization of ASIC1a is likely important for its regulation of spine morphology and synaptic functions. Our findings reveal novel insights into the mechanism that governs ASIC1a expression and function at peri-synaptic regions of dendritic spines, which may be targeted to manipulate synaptic dysfunction in disease therapies. Along with the knowledge that both PS and ASIC1a are crucial for the normal function of neurons, it will be interesting to test their involvement in neurodevelopment and neurological diseases, such as lipid disorders with neurological complications.

## Acknowledgements

This study was supported by grants from the National Natural Science Foundation of China (81961128024, 81730095, 32170994), US National Institutes of Health (NS1147167), the Shanghai Municipal Science and Technology Major Project (2018SHZDZX05) and Innovative Research Team of High-level Local Universities in Shanghai. We thank Prof. Michael J. Welsh (Howard Hughes Medical Institute, University of Iowa, USA) for providing *Asic1a^-/-^*mice; Dr. Bo Duan, Dr. Wei-Zheng Zeng, Ms. Ying Li, Ms. Hui Cao, and Ms. Jin Cheng for technical assistance; and Prof. Nan-Jie Xu (Department of Anatomy and Physiology, Shanghai Jiao Tong University School of Medicine, China) for providing reagent and helpful comments on this manuscript. We also thank Ms. Ying Shi (Institute of Brain Science, Fudan University), Ms. Yan Wang, Ms. Yang Yu, and Ms. Na Liu (National Center for Protein Science, Shanghai) for kind help with the SIM imaging work.

## References

1. J. G. Kay, S. Grinstein, Phosphatidylserine-mediated cellular signaling. Adv Exp Med Biol 991, 177–193 (2013).

2. J. A. Allen, R. A. Halverson-Tamboli, M. M. Rasenick, Lipid raft microdomains and neurotransmitter signalling. Nat Rev Neurosci 8, 128-140 (2007).

3. K. L. Arendt, M. Royo, M. Fernandez-Monreal, S. Knafo, C. N. Petrok, J. R. Martens, J. A. Esteban, PIP3 controls synaptic function by maintaining AMPA receptor clustering at the postsynaptic membrane. Nat Neurosci 13, 36–44 (2010).

4. T. M. Khuong, R. L. Habets, S. Kuenen, A. Witkowska, J. Kasprowicz, J. Swerts, R. Jahn, G. van den Bogaart, P. Verstreken, Synaptic PI(3,4,5)P3 is required for Syntaxin1A clustering and neurotransmitter release. Neuron 77, 1097–1108 (2013).

5. G. D. Fairn, M. Hermansson, P. Somerharju, S. Grinstein, Phosphatidylserine is polarized and required for proper Cdc42 localization and for development of cell polarity. Nature cell biology 13, 1424-1430 (2011).

6. M. Tsuchiya, Y. Hara, M. Okuda, K. Itoh, R. Nishioka, A. Shiomi, K. Nagao, M. Mori, Y. Mori, J. Ikenouchi, R. Suzuki, M. Tanaka, T. Ohwada, J. Aoki, M. Kanagawa, T. Toda, Y. Nagata, R. Matsuda, Y. Takayama, M. Tominaga, M. Umeda, Cell surface flip-flop of phosphatidylserine is critical for PIEZO1-mediated myotube formation. Nature communications 9, 2049 (2018).

7. D. Bach, R. F. Epand, R. M. Epand, E. Wachtel, Interaction of 7-ketocholesterol with two major components of the inner leaflet of the plasma membrane: phosphatidylethanolamine and phosphatidylserine. Biochemistry (Mosc). 47, 3004-3012 (2008).

8. T. Yeung, G. E. Gilbert, J. Shi, J. Silvius, A. Kapus, S. Grinstein, Membrane phosphatidylserine regulates surface charge and protein localization. Science 319, 210-213 (2008).

9. R. Raghupathy, A. A. Anilkumar, A. Polley, P. P. Singh, M. Yadav, C. Johnson, S. Suryawanshi, V. Saikam, S. D. Sawant, A. Panda, Z. Guo, R. A. Vishwakarma, M. Rao, S. Mayor, Transbilayer lipid interactions mediate nanoclustering of lipid-anchored proteins. Cell 161, 581-594 (2015).

10. J. T. Shieh, S. Aradhya, A. Novelli, M. A. Manning, A. M. Cherry, J. Brumblay, C. D. Salpietro, L. Bernardini, B. Dallapiccola, H. E. Hoyme, Nablus mask-like facial syndrome is caused by a microdeletion of 8q detected by array-based comparative genomic hybridization. Am J Med Genet A 140, 1267-1273 (2006).

11. S. B. Sousa, D. Jenkins, E. Chanudet, G. Tasseva, M. Ishida, G. Anderson, J. Docker, M. Ryten, J. Sa, J. M. Saraiva, A. Barnicoat, R. Scott, A. Calder, D. Wattanasirichaigoon, K. Chrzanowska, M. Simandlova, L. Van Maldergem, P. Stanier, P. L. Beales, J. E. Vance, G. E. Moore, Gain-of-function mutations in the phosphatidylserine synthase 1 (PTDSS1) gene cause Lenz-Majewski syndrome. Nat Genet 46, 70-76 (2013).

12. N. Scott-Hewitt, F. Perrucci, R. Morini, M. Erreni, M. Mahoney, A. Witkowska, A. Carey, E. Faggiani, L. T. Schuetz, S. Mason, M. Tamborini, M. Bizzotto, L. Passoni, F. Filipello, R. Jahn, B. Stevens, M. Matteoli, Local externalization of phosphatidylserine mediates developmental synaptic pruning by microglia. EMBO J. 39, e105380 (2020).

13. T. Li, B. Chiou, C. K. Gilman, R. Luo, T. Koshi, D. Yu, H. C. Oak, S. Giera, E. Johnson-Venkatesh, A. K. Muthukumar, B. Stevens, H. Umemori, X. Piao, A splicing isoform of GPR56 mediates microglial synaptic refinement via phosphatidylserine binding. EMBO J. 39, e104136 (2020).

14. D. Sokolova, T. Childs, S. Hong, Insight into the role of phosphatidylserine in complement-mediated synapse loss in Alzheimer’s disease. Fac Rev 10, 19 (2021).

15. J. Garcia-Anoveros, B. Derfler, J. Neville-Golden, B. T. Hyman, D. P. Corey, BNaC1 and BNaC2 constitute a new family of human neuronal sodium channels related to degenerins and epithelial sodium channels. Proc. Natl. Acad. Sci. U. S. A. 94, 1459-1464 (1997).

16. R. Waldmann, G. Champigny, F. Bassilana, C. Heurteaux, M. Lazdunski, A proton-gated cation channel involved in acid-sensing. Nature 386, 173-177 (1997).

17. T. W. Sherwood, K. G. Lee, M. G. Gormley, C. C. Askwith, Heteromeric acid-sensing ion channels (ASICs) composed of ASIC2b and ASIC1a display novel channel properties and contribute to acidosis-induced neuronal death. J Neurosci 31, 9723-9734 (2011).

18. X. L. Song, D. S. Liu, M. Qiang, Q. Li, M. G. Liu, W. G. Li, X. Qi, N. J. Xu, G. Yang, M. X. Zhu, T. L. Xu, Postsynaptic Targeting and Mobility of Membrane Surface-Localized hASIC1a. Neurosci Bull 37, 145-165 (2021).

19. X. M. Zha, J. A. Wemmie, S. H. Green, M. J. Welsh, Acid-sensing ion channel 1a is a postsynaptic proton receptor that affects the density of dendritic spines. Proc Natl Acad Sci U S A 103, 16556–16561 (2006).

20. X. M. Zha, V. Costa, A. M. Harding, L. Reznikov, C. J. Benson, M. J. Welsh, ASIC2 subunits target acid-sensing ion channels to the synapse via an association with PSD-95. J Neurosci 29, 8438–8446 (2009).

21. C. Gonzalez-Inchauspe, F. J. Urbano, M. N. Di Guilmi, O. D. Uchitel, Acid sensing ion channels activated by evoked released protons modulate synaptic transmission at the mouse calyx of Held synapse. J Neurosci 37, 2589-2599 (2017).

22. C. J. Kreple, Y. Lu, R. J. Taugher, A. L. Schwager-Gutman, J. Du, M. Stump, Y. Wang, A. Ghobbeh, R. Fan, C. V. Cosme, L. P. Sowers, M. J. Welsh, J. J. Radley, R. T. LaLumiere, J. A. Wemmie, Acid-sensing ion channels contribute to synaptic transmission and inhibit cocaine-evoked plasticity. Nat Neurosci 17, 1083-1091 (2014).

23. J. A. Wemmie, J. Chen, C. C. Askwith, A. M. Hruska-Hageman, M. P. Price, B. C. Nolan, P. G. Yoder, E. Lamani, T. Hoshi, J. H. Freeman, Jr., M. J. Welsh, The acid-activated ion channel ASIC contributes to synaptic plasticity, learning, and memory. Neuron 34, 463-477 (2002).

24. P. H. Chiang, T. C. Chien, C. C. Chen, Y. Yanagawa, C. C. Lien, ASIC-dependent LTP at multiple glutamatergic synapses in amygdala network is required for fear memory. Scientific reports 5, 10143 (2015).

25. M. G. Liu, H. S. Li, W. G. Li, Y. J. Wu, S. N. Deng, C. Huang, O. Maximyuk, V. Sukach, O. Krishtal, M. X. Zhu, T. L. Xu, Acid-sensing ion channel 1a contributes to hippocampal LTP inducibility through multiple mechanisms. Scientific reports 6, 23350 (2016).

26. S. Chai, M. Li, D. Branigan, Z. G. Xiong, R. P. Simon, Activation of acid-sensing ion channel 1a (ASIC1a) by surface trafficking. J. Biol. Chem. 285, 13002-13011 (2010).

27. W. Z. Zeng, D. S. Liu, B. Duan, X. L. Song, X. Wang, D. Wei, W. Jiang, M. X. Zhu, Y. Li, T. L. Xu, Molecular Mechanism of Constitutive Endocytosis of Acid-Sensing Ion Channel 1a and Its Protective Function in Acidosis-Induced Neuronal Death. J Neurosci 33, 7066-7078 (2013).

28. B. Duan, D. S. Liu, Y. Huang, W. Z. Zeng, X. Wang, H. Yu, M. X. Zhu, Z. Y. Chen, T. L. Xu, PI3-kinase/Akt pathway-regulated membrane insertion of acid-sensing ion channel 1a underlies BDNF-induced pain hypersensitivity. J. Neurosci. 32, 6351-6363 (2012).

29. Q. F. Wu, L. Yang, S. Li, Q. Wang, X. B. Yuan, X. Gao, L. Bao, X. Zhang, Fibroblast growth factor 13 is a microtubule-stabilizing protein regulating neuronal polarization and migration. Cell 149, 1549-1564 (2012).

30. T. Saito, N. Nakatsuji, Efficient gene transfer into the embryonic mouse brain using in vivo electroporation. Dev Biol 240, 237-246 (2001).

31. L. J. Wu, B. Duan, Y. D. Mei, J. Gao, J. G. Chen, M. Zhuo, L. Xu, M. Wu, T. L. Xu, Characterization of acid-sensing ion channels in dorsal horn neurons of rat spinal cord. J Biol Chem 279, 43716–43724 (2004).

32. S. J. Martin, C. P. Reutelingsperger, A. J. McGahon, J. A. Rader, R. C. van Schie, D. M. LaFace, D. R. Green, Early redistribution of plasma membrane phosphatidylserine is a general feature of apoptosis regardless of the initiating stimulus: inhibition by overexpression of Bcl-2 and Abl. J Exp Med 182, 1545-1556 (1995).

33. A. L. Mammen, R. L. Huganir, R. J. O’Brien, Redistribution and stabilization of cell surface glutamate receptors during synapse formation. J Neurosci 17, 7351-7358 (1997).

34. W. J. Tyler, L. Pozzo-Miller, Miniature synaptic transmission and BDNF modulate dendritic spine growth and form in rat CA1 neurones. J Physiol 553, 497-509 (2003).

35. B. Duan, L. J. Wu, Y. Q. Yu, Y. Ding, L. Jing, L. Xu, J. Chen, T. L. Xu, Upregulation of acid-sensing ion channel ASIC1a in spinal dorsal horn neurons contributes to inflammatory pain hypersensitivity. J Neurosci 27, 11139–11148 (2007).

36. M. Simons, W. J. Gault, D. Gotthardt, R. Rohatgi, T. J. Klein, Y. Shao, H. J. Lee, A. L. Wu, Y. Fang, L. M. Satlin, J. T. Dow, J. Chen, J. Zheng, M. Boutros, M. Mlodzik, Electrochemical cues regulate assembly of the Frizzled/Dishevelled complex at the plasma membrane during planar epithelial polarization. Nat Cell Biol 11, 286–294 (2009).

37. T. Freisinger, R. Wedlich-Soldner, Phosphatidylserine promotes polar Cdc42 localization. Nat Cell Biol 13, 1387-1388 (2011).

38. J. G. Kay, M. Koivusalo, X. Ma, T. Wohland, S. Grinstein, Phosphatidylserine dynamics in cellular membranes. Mol Biol Cell 23, 2198-2212 (2012).

39. T. M. Newpher, M. D. Ehlers, Glutamate receptor dynamics in dendritic microdomains. Neuron 58, 472-497 (2008).

40. F. Calderon, H. Y. Kim, Detection of intracellular phosphatidylserine in living cells. J Neurochem 104, 1271-1279 (2008).

41. D. Alvarez de la Rosa, S. R. Krueger, A. Kolar, D. Shao, R. M. Fitzsimonds, C. M. Canessa, Distribution, subcellular localization and ontogeny of ASIC1 in the mammalian central nervous system. J Physiol 546, 77-87 (2003).

42. R. Mozzi, S. Buratta, G. Goracci, Metabolism and functions of phosphatidylserine in mammalian brain. Neurochem Res 28, 195-214 (2003).

43. R. Mozzi, S. Buratta, “Brain Phosphatidylserine: Metabolism and Functions” in Handbook of Neurochemistry and Molecular Neurobiology, A. Lajtha, G. Tettamanti, G. Goracci, Eds. (Springer US, 2010), chap. 3, pp. 39–58.

44. J. A. Wemmie, C. C. Askwith, E. Lamani, M. D. Cassell, J. H. Freeman, Jr., M. J. Welsh, Acid-sensing ion channel 1 is localized in brain regions with high synaptic density and contributes to fear conditioning. J Neurosci 23, 5496-5502 (2003).

45. L. Jing, X. P. Chu, Y. Q. Jiang, D. M. Collier, B. Wang, Q. Jiang, P. M. Snyder, X. M. Zha, N-glycosylation of acid-sensing ion channel 1a regulates its trafficking and acidosis-induced spine remodeling. J Neurosci 32, 4080-4091 (2012).

46. O. Kuge, M. Nishijima, Phosphatidylserine synthase I and II of mammalian cells. Biochim Biophys Acta 1348, 151-156 (1997).

47. O. Kuge, K. Saito, M. Nishijima, Cloning of a Chinese hamster ovary (CHO) cDNA encoding phosphatidylserine synthase (PSS) II, overexpression of which suppresses the phosphatidylserine biosynthetic defect of a PSS I-lacking mutant of CHO-K1 cells. J Biol Chem 272, 19133-19139 (1997).

48. S. J. Stone, J. E. Vance, Phosphatidylserine synthase-1 and -2 are localized to mitochondria-associated membranes. J Biol Chem 275, 34534-34540 (2000).

49. J. E. Vance, G. Tasseva, Formation and function of phosphatidylserine and phosphatidylethanolamine in mammalian cells. Biochim Biophys Acta 1831, 543-554 (2013).

50. O. Kuge, M. Nishijima, Y. Akamatsu, Phosphatidylserine biosynthesis in cultured Chinese hamster ovary cells. II. Isolation and characterization of phosphatidylserine auxotrophs. J Biol Chem 261, 5790-5794 (1986).

51. O. Kuge, M. Nishijima, Y. Akamatsu, A Chinese hamster cDNA encoding a protein essential for phosphatidylserine synthase I activity. J Biol Chem 266, 24184-24189 (1991).

52. I. Vermes, C. Haanen, H. Steffens-Nakken, C. Reutelingsperger, A novel assay for apoptosis. Flow cytometric detection of phosphatidylserine expression on early apoptotic cells using fluorescein labelled Annexin V. J Immunol Methods 184, 39-51 (1995).

53. M. Akbar, F. Calderon, Z. Wen, H. Y. Kim, Docosahexaenoic acid: a positive modulator of Akt signaling in neuronal survival. Proc Natl Acad Sci U S A 102, 10858-10863 (2005).

54. M. A. Lemmon, Membrane recognition by phospholipid-binding domains. Nat Rev Mol Cell Biol 9, 99-111 (2008).

55. G. Di Paolo, P. De Camilli, Phosphoinositides in cell regulation and membrane dynamics. Nature 443, 651-657 (2006).

56. T. Rohacs, C. M. Lopes, I. Michailidis, D. E. Logothetis, PI(4,5)P2 regulates the activation and desensitization of TRPM8 channels through the TRP domain. Nat. Neurosci. 8, 626-634 (2005).

57. X. P. Dong, D. Shen, X. Wang, T. Dawson, X. Li, Q. Zhang, X. Cheng, Y. Zhang, L. S. Weisman, M. Delling, H. Xu, PI(3,5)P(2) controls membrane trafficking by direct activation of mucolipin Ca(2+) release channels in the endolysosome. Nat Commun 1, 38 (2010).

58. L. P. Bernier, D. Blais, E. Boue-Grabot, P. Seguela, A dual polybasic motif determines phosphoinositide binding and regulation in the P2X channel family. PLoS One 7, e40595 (2012).

59. L. P. Bernier, A. R. Ase, S. Chevallier, D. Blais, Q. Zhao, E. Boue-Grabot, D. Logothetis, P. Seguela, Phosphoinositides regulate P2X4 ATP-gated channels through direct interactions. J Neurosci 28, 12938-12945 (2008).

60. J. A. Saugstad, J. A. Roberts, J. Dong, S. Zeitouni, R. J. Evans, Analysis of the membrane topology of the acid-sensing ion channel 2a. J. Biol. Chem. 279, 55514-55519 (2004).

61. N. Yoder, C. Yoshioka, E. Gouaux, Gating mechanisms of acid-sensing ion channels. Nature 555, 397-401 (2018).

62. H. Hering, C. C. Lin, M. Sheng, Lipid rafts in the maintenance of synapses, dendritic spines, and surface AMPA receptor stability. J Neurosci 23, 3262-3271 (2003).

63. A. Hall, Rho GTPases and the actin cytoskeleton. Science 279, 509-514 (1998).

64. Y. Z. Wang, W. Z. Zeng, X. Xiao, Y. Huang, X. L. Song, Z. Yu, D. Tang, X. P. Dong, M. X. Zhu, T. L. Xu, Intracellular ASIC1a regulates mitochondrial permeability transition-dependent neuronal death. Cell Death Differ 20, 1359-1369 (2013).

65. G. D. Fairn, N. L. Schieber, N. Ariotti, S. Murphy, L. Kuerschner, R. I. Webb, S. Grinstein, R. G. Parton, High-resolution mapping reveals topologically distinct cellular pools of phosphatidylserine. J Cell Biol 194, 257-275 (2011).

66. W. Chen, Y. Feng, D. Chen, A. Wandinger-Ness, Rab11 is required for trans-golgi network-to-plasma membrane transport and a preferential target for GDP dissociation inhibitor. Mol Biol Cell 9, 3241-3257 (1998).

67. R. Sannerud, J. Saraste, B. Goud, Retrograde traffic in the biosynthetic-secretory route: pathways and machinery. Curr Opin Cell Biol 15, 438-445 (2003).

68. P. A. Leventis, S. Grinstein, “The Distribution and Function of Phosphatidylserine in Cellular Membranes” in Annual Review of Biophysics, Vol 39, D. C. Rees, K. A. Dill, J. R. Williamson, Eds. (Annual Review of Biophysics, 2010), vol. 39, pp. 407–427.

69. Y. Uchida, J. Hasegawa, D. Chinnapen, T. Inoue, S. Okazaki, R. Kato, S. Wakatsuki, R. Misaki, M. Koike, Y. Uchiyama, S. Iemura, T. Natsume, R. Kuwahara, T. Nakagawa, K. Nishikawa, K. Mukai, E. Miyoshi, N. Taniguchi, D. Sheff, W. I. Lencer, T. Taguchi, H. Arai, Intracellular phosphatidylserine is essential for retrograde membrane traffic through endosomes. Proc Natl Acad Sci U S A 108, 15846-15851 (2011).

70. J. F. Hancock, K. Cadwallader, H. Paterson, C. J. Marshall, A CAAX or a CAAL motif and a second signal are sufficient for plasma membrane targeting of ras proteins. EMBO J 10, 4033-4039 (1991).

71. Y. Posor, M. Eichhorn-Gruenig, D. Puchkov, J. Schoneberg, A. Ullrich, A. Lampe, R. Muller, S. Zarbakhsh, F. Gulluni, E. Hirsch, M. Krauss, C. Schultz, J. Schmoranzer, F. Noe, V. Haucke, Spatiotemporal control of endocytosis by phosphatidylinositol-3,4-bisphosphate. Nature 499, 233-237 (2013).

72. F. Wang, P. Herzmark, O. D. Weiner, S. Srinivasan, G. Servant, H. R. Bourne, Lipid products of PI(3)Ks maintain persistent cell polarity and directed motility in neutrophils. Nat Cell Biol 4, 513-518 (2002).

73. N. Lucas, W. Cho, Phosphatidylserine binding is essential for plasma membrane recruitment and signaling function of 3-phosphoinositide-dependent kinase-1. J Biol Chem 286, 41265-41272 (2011).

74. J. N. Mazerik, M. J. Tyska, Myosin-1A targets to microvilli using multiple membrane binding motifs in the tail homology 1 (TH1) domain. J Biol Chem 287, 13104-13115 (2012).

75. Z. C. Abay, M. Y.-Y. Wong, J.-S. Teoh, T. Vijayaraghavan, M. A. Hilliard, B. Neumann, Phosphatidylserine save-me signals drive functional recovery of severed axons in Caenorhabditis elegans. Proc Natl Acad Sci U S A 114, E10196-E10205 (2017).

76. G. Di Paolo, H. S. Moskowitz, K. Gipson, M. R. Wenk, S. Voronov, M. Obayashi, R. Flavell, R. M. Fitzsimonds, T. A. Ryan, P. De Camilli, Impaired PtdIns(4,5)P2 synthesis in nerve terminals produces defects in synaptic vesicle trafficking. Nature 431, 415-422 (2004).

77. K. Schwarz, S. Natarajan, N. Kassas, N. Vitale, F. Schmitz, The synaptic ribbon is a site of phosphatidic acid generation in ribbon synapses. J Neurosci 31, 15996-16011 (2011).

78. L. Trovo, T. Ahmed, Z. Callaerts-Vegh, A. Buzzi, C. Bagni, M. Chuah, T. Vandendriessche, R. D’Hooge, D. Balschun, C. G. Dotti, Low hippocampal PI(4,5)P(2) contributes to reduced cognition in old mice as a result of loss of MARCKS. Nat Neurosci 16, 449-455 (2013).

79. A. P. H. de Jong, C. M. Roggero, M. R. Ho, M. Y. Wong, C. A. Brautigam, J. Rizo, P. S. Kaeser, RIM C2B Domains Target Presynaptic Active Zone Functions to PIP2-Containing Membranes. Neuron 98, 335-349 e337 (2018).

80. E. Sartorel, C. Unlu, M. Jose, A. Massoni-Laporte, J. Meca, J. B. Sibarita, D. McCusker, Phosphatidylserine and GTPase activation control Cdc42 nanoclustering to counter dissipative diffusion. Mol Biol Cell 29, 1299-1310 (2018).

81. A. C. Newton, D. E. Koshland, Jr., Phosphatidylserine affects specificity of protein kinase C substrate phosphorylation and autophosphorylation. Biochemistry 29, 6656-6661 (1990).

82. S. Ghosh, W. Q. Xie, A. F. Quest, G. M. Mabrouk, J. C. Strum, R. M. Bell, The cysteine-rich region of raf-1 kinase contains zinc, translocates to liposomes, and is adjacent to a segment that binds GTP-ras. J Biol Chem 269, 10000-10007 (1994).

83. T. Yeung, M. Terebiznik, L. Yu, J. Silvius, W. M. Abidi, M. Philips, T. Levine, A. Kapus, S. Grinstein, Receptor activation alters inner surface potential during phagocytosis. Science 313, 347-351 (2006).

84. Y. Zhou, C. O. Wong, K. J. Cho, D. van der Hoeven, H. Liang, D. P. Thakur, J. Luo, M. Babic, K. E. Zinsmaier, M. X. Zhu, H. Hu, K. Venkatachalam, J. F. Hancock, SIGNAL TRANSDUCTION. Membrane potential modulates plasma membrane phospholipid dynamics and K-Ras signaling. Science 349, 873-876 (2015).

85. E. Hildebrandt, N. Khazanov, J. C. Kappes, Q. Dai, H. Senderowitz, I. L. Urbatsch, Specific stabilization of CFTR by phosphatidylserine. Biochim Biophys Acta 1859, 289-293 (2017).

86. S. Marra, R. Ferru-Clement, V. Breuil, A. Delaunay, M. Christin, V. Friend, S. Sebille, C. Cognard, T. Ferreira, C. Roux, L. Euller-Ziegler, J. Noel, E. Lingueglia, E. Deval, Non-acidic activation of pain-related Acid-Sensing Ion Channel 3 by lipids. EMBO J. 35, 414-428 (2016).

87. E. S. Smith, H. Cadiou, P. A. McNaughton, Arachidonic acid potentiates acid-sensing ion channels in rat sensory neurons by a direct action. Neuroscience 145, 686-698 (2007).

88. Z. Yu, Y. J. Wu, Y. Z. Wang, D. S. Liu, X. L. Song, Q. Jiang, Y. Li, S. Zhang, N. J. Xu, M. X. Zhu, W. G. Li, T. L. Xu, The acid-sensing ion channel ASIC1a mediates striatal synapse remodeling and procedural motor learning. Sci Signal 11, pii: eaar4481 (2018).

89. H. S. Li, X. Y. Su, X. L. Song, X. Qi, Y. Li, R. Q. Wang, O. Maximyuk, O. Krishtal, T. Wang, H. Fang, L. Liao, H. Cao, Y. Q. Zhang, M. X. Zhu, M. G. Liu, T. L. Xu, Protein Kinase C Lambda Mediates Acid-Sensing Ion Channel 1a-Dependent Cortical Synaptic Plasticity and Pain Hypersensitivity. J. Neurosci. 39, 5773-5793 (2019).

90. A. E. Ziemann, J. E. Allen, N. S. Dahdaleh, Drebot, II, M. W. Coryell, A. M. Wunsch, C. M. Lynch, F. M. Faraci, M. A. Howard, 3rd, M. J. Welsh, J. A. Wemmie, The amygdala is a chemosensor that detects carbon dioxide and acidosis to elicit fear behavior. Cell 139, 1012-1021 (2009).

91. H. Y. Kim, B. X. Huang, A. A. Spector, Phosphatidylserine in the brain: metabolism and function. Prog Lipid Res 56, 1-18 (2014).

92. Q. L. Ma, X. Zuo, F. Yang, O. J. Ubeda, D. J. Gant, M. Alaverdyan, N. C. Kiosea, S. Nazari, P. P. Chen, F. Nothias, P. Chan, E. Teng, S. A. Frautschy, G. M. Cole, Loss of MAP function leads to hippocampal synapse loss and deficits in the Morris Water Maze with aging. J Neurosci 34, 7124-7136 (2014).

93. D. Arikketh, R. Nelson, J. E. Vance, Defining the importance of phosphatidylserine synthase-1 (PSS1): unexpected viability of PSS1-deficient mice. J Biol Chem 283, 12888-12897 (2008).

94. A. Baude, Z. Nusser, J. D. Roberts, E. Mulvihill, R. A. McIlhinney, P. Somogyi, The metabotropic glutamate receptor (mGluR1 alpha) is concentrated at perisynaptic membrane of neuronal subpopulations as detected by immunogold reaction. Neuron 11, 771-787 (1993).

95. A. M. Harding, N. Kusama, T. Hattori, M. Gautam, C. J. Benson, ASIC2 Subunits Facilitate Expression at the Cell Surface and Confer Regulation by PSD-95. PLoS One 9, e93797 (2014).

96. M. Li, E. Kratzer, K. Inoue, R. P. Simon, Z. G. Xiong, Developmental change in the electrophysiological and pharmacological properties of acid-sensing ion channels in CNS neurons. J Physiol 588, 3883-3900 (2010).

97. W. G. Li, M. G. Liu, S. Deng, Y. M. Liu, L. Shang, J. Ding, T. T. Hsu, Q. Jiang, Y. Li, F. Li, M. X. Zhu, T. L. Xu, ASIC1a regulates insular long-term depression and is required for the extinction of conditioned taste aversion. Nature communications 7, 13770 (2016).

98. S. Sankaranarayanan, T. A. Ryan, Real-time measurements of vesicle-SNARE recycling in synapses of the central nervous system. Nat Cell Biol 2, 197-204 (2000).

99. H. P. Ma, S. Saxena, D. G. Warnock, Anionic phospholipids regulate native and expressed epithelial sodium channel (ENaC). J Biol Chem 277, 7641-7644 (2002).

100. Z. R. Zhang, C. F. Chou, J. Wang, Y. Y. Liang, H. P. Ma, Anionic phospholipids differentially regulate the epithelial sodium channel (ENaC) by interacting with alpha, beta, and gamma ENaC subunits. Pflugers Arch 459, 377-387 (2010).

101. J. Jasti, H. Furukawa, E. B. Gonzales, E. Gouaux, Structure of acid-sensing ion channel 1 at 1.9 A resolution and low pH. Nature 449, 316-323 (2007).

102. E. B. Gonzales, T. Kawate, E. Gouaux, Pore architecture and ion sites in acid-sensing ion channels and P2X receptors. Nature 460, 599-604 (2009).

103. O. Cremona, G. Di Paolo, M. R. Wenk, A. Luthi, W. T. Kim, K. Takei, L. Daniell, Y. Nemoto, S. B. Shears, R. A. Flavell, D. A. McCormick, P. De Camilli, Essential role of phosphoinositide metabolism in synaptic vesicle recycling. Cell 99, 179-188 (1999).

104. P. Xu, R. D. Baldridge, R. J. Chi, C. G. Burd, T. R. Graham, Phosphatidylserine flipping enhances membrane curvature and negative charge required for vesicular transport. J Cell Biol 202, 875-886 (2013).

105. H. T. McMahon, J. L. Gallop, Membrane curvature and mechanisms of dynamic cell membrane remodelling. Nature 438, 590-596 (2005).

106. T. Matsudaira, K. Mukai, T. Noguchi, J. Hasegawa, T. Hatta, S. I. Iemura, T. Natsume, N. Miyamura, H. Nishina, J. Nakayama, K. Semba, T. Tomita, S. Murata, H. Arai, T. Taguchi, Endosomal phosphatidylserine is critical for the YAP signalling pathway in proliferating cells. Nat Commun 8, 1246 (2017).

