## Supplemental information for "Phosphatidylserine controls synaptic targeting and membrane stability of ASIC1a"

**Figure S1. Two other examples of phospholipid-strip assay. Supplementary information for Figure 1. A.** GST-$Ct_{ASIC1a}$ fusion proteins were incubated with the membrane strips under different buffering systems: PBS buffer (left, 140 mM NaCl, 3 mM KCl, 10 mM phosphate buffer, 0.1% Tween-20, pH 7.4), or TBS buffer (right, 10 mM Tris, 150 mM NaCl, 0.1% Tween-20, pH 8.0), and then probed with anti-GST antibody. **B.** Detection of purified GST, GST-$Nt_{ASIC1a}$, GST-$Ct_{ASIC1a}$, GST-$Nt_{ASIC2a}$, and GST-$Ct_{ASIC2a}$ fusion proteins by Coomassie Blue staining and Western Blotting. **C.** Phospholipid-strip detection of GST proteins fused with the intracellular C-terminal domains of glutamate receptors GluA1, GluA2 and GluN1. **D.** Detection of purified GST, GST-$Ct_{GluA1}$, GST-$Ct_{GluA2}$, GST-$Ct_{GluN1}$ fusion proteins by Coomassie Blue staining and Western Blotting. **E.** Diagram illustration of membrane topology and cytosolic domains (orange) of ASIC1a, ASIC2a, GluA1, GluA2 and GluN1.

**Figure S2. Spine enrichment and synaptic location of ASIC1a and Lact-C2. Supplementary information for Figure 2A-2C.**

**A.** Representative confocal images of DIV21-28 rat cortical neurons transfected with EGFP and mCherry-Lact-C2 (left), EGFP and mCherry-ASIC1a (middle), and EGFP and mCherry (right) constructs at DIV7. After fixation, neurons were immunostained with the antibodies of corresponding fluorescent proteins. Examples for neuronal dendrites (blue rectangle) are also shown in high-magnification images. **B.** Representative confocal images of DIV21-28 cortical neurons transfected with EGFP and mCherry-Lact-C2-AAA at DIV7. Examples for neuronal dendrites (blue rectangle) are also shown in high-magnification images. **C.** Comparison of the distributions between wild type Lact-C2 and Lact-C2-AAA in cortical neurons. Signal variability of each group is indicated by coefficient of variation (C.V.). n = 10 cells. ***$p$ < 0.001 (unpaired Student's $t$-test). **D.** Comparison of the distributions between Lact-C2 and Lact-C2-AAA in different compartments of DIV21-28 rat cortical neurons. n = 10 cells. **$p$ = 0.0075, ***$p$ < 0.001 (unpaired Student's $t$-test). **E.** Representative confocal images of DIV28 rat cortical neurons transfected with EGFP at DIV7. After fixation, the neurons were immunostained with the antibodies of PSD-95 (left) or

Synapsin I (right).

**Figure S3. Super-resolution images showing synaptic localization of ASIC1a and Lact-C2. Supplementary information for Figure 2.**

**A.** Representative SIM z-stack maximum projection images of one dendrite of DIV28 rat cortical neuron transfected with EGFP-ASIC1a and mCherry-Lact-C2 on DIV7, imaged with structured illumination microscopy (SIM). After fixation, neurons were immunostained with the antibodies of PSD-95 and corresponding fluorescent proteins. Red arrowheads indicate PSD areas. Spines marked with numbers (in the lower panel) are shown sequentially in (B). **B.** Six examples of spines in the same dendrite shown in (A). Arrowheads, PSD areas; white lines, line scan areas in (C). **C.** Intensity profiling of line scan areas through the spine heads from (B). ASIC1a, green lines; Lact-C2, red lines; PSD-95, blue lines. **D.** 3D reconstruction of the six example spines in (B). Arrowheads, PSD areas. **E.** Correlation plots of the areas of spine ASIC1a (green, $R^2$ = 0.7547, slope = 0.5066 ± 0.0222, $p$ < 0.0001, Spearman's test) or Lact-C2 (red, $R^2$ = 0.7781, slope = 0.5674 ± 0.0232, $p$ < 0.0001, Spearman's test) vs. spine head areas, n = 172 spines from 7 cells. **F.** Correlation plots of the areas of spine ASIC1a (green, $R^2$ = 0.5226, slope = 1.335 ± 0.0979, $p$ < 0.0001, Spearman's test) or Lact-C2 (red, $R^2$ = 0.4753, slope = 1.405 ± 0.1132, $p$ < 0.0001, Spearman's test) vs. PSD areas, n = 172 spines from 7 cells. **G.** Correlation plots of PSD areas vs. spine head areas ($R^2$ = 0.4850, slope = 0.2199 ± 0.0174, $p$ < 0.0001, Spearman's test), n = 172 spines from 7 cells. **H**. Representative SIM z-stack maximum projection image (left) and 3D reconstruction (right) of one dendrite of DIV28 *Asic1a*$^{-/-}$ mouse cortical neuron transfected with HA-ASIC1a on DIV7. Neurons were stained with anti-HA antibody under non-permeabilized conditions for surface ASIC1a and with anti-PSD-95 antibody under permeabilized conditions for PSD-95. **I.** Cumulative probabilities of densities of surface ASIC1a nanoclusters distributed in different compartments of cortical neurons. **J & K.** Frequency distributions of the numbers of surface ASIC1a nanoclusters in spine head (J) or spine neck (K). **L.** Frequency distribution of the diameters (semi-major axis) of surface ASIC1a nanoclusters.

**Figure S4. Postsynaptic localizations of GluN1.**

**A.** Representative super-resolution images of a DIV28 rat cortical neuron transfected with EGFP-GluN1 and mCherry on DIV7, imaged with SIM. After fixation, neurons were immunostained with the antibodies of PSD-95 and corresponding fluorescent protein. More details about the spines marked with numbers are shown individually in (B) and (C). **B.** SIM z stack maximum projection images of dendritic spines marked in (A). White lines, line scan areas through the spine heads, with the intensity profiling shown at right. Green lines, GluN1; red dash lines, mCherry; blue lines, PSD-95. **C.** 3D reconstruction images of the spine heads shown in (B). **D.** Colocalization analysis between GluN1 and PSD-95 (or mCherry). Bars indicate mean ± SEM, n = 50 spines from 10 cells. ***$p < 0.001$ (Student's *t*-test). **E.** Schematic diagram of four patterns of synaptic localizations of GluN1 and PSD-95. Green, GluN1; blue, PSD-95; cyan, colocalization area.

**Figure S5. Comparison of confocal and Airyscan imaging results.**

**A.** Representative confocal image and super-resolution image (Airyscan) of the same DIV28 neuron transfected with EGFP-ASIC1a and mCherry-Lact-C2 on DIV7. After fixation, neurons were immunostained with the antibodies of PSD-95 and corresponding fluorescent proteins. **B.** Comparison between confocal and Airyscan images of the same neuronal dendrite. Scale bar = 5 μm. **C-E.** Colocalization analysis between ASIC1a and Lact-C2 (C), Lact-C2 and PSD-95 (D), ASIC1a and PSD-95 (E). Bars indicate mean ± SEM, n = 9 cells. *$p = 0.0242$, ***$p = 0.0004$ (Student's *t*-test).

**Figure S6**. **Surface distribution of ASIC1a in cortical neurons.**

**A & B.** Representative confocal images of DIV28 cultured rat cortical neurons transfected with HA-ASIC1a on DIV7. After fixation, neurons were immunostained before permeabilization with monoclonal HA antibody followed by Alexa-488-labeled secondary antibody (1$^{st}$ panel from left) to visualize surface ASIC1a and after permeabilization with monoclonal HA antibody (A) or polyclonal HA antibody (B) followed by corresponding Alexa-568-labeled secondary antibody (2nd panel) to label total or cytoplasmic ASIC1a, respectively. 3rd panel, merged image for 1st and 2nd panels; 4th panel, another example in the same experiment. C, cytoplasmic ASIC1a; S, surface ASIC1a; T, total ASIC1a. **C.** High-magnification images of neuronal dendrites in boxed areas of (A) and (B). **D.** Surface fraction of ASIC1a (surface/total ratio) during neurodevelopment *in vitro* (2-5 weeks). n = 5. $p$ = 0.0007, vs. DIV14 group (one-way ANOVA with Tukey's post hoc); $*p < 0.05$, $**p < 0.01$, $***p < 0.001$.

**Figure S7**. **Juxtamembranous location of PS-binding motif.**

**A.** Sequences and structures of TMII and proximal C-terminus of ASIC1a. **A1**, Alignment of juxtamembranous domains of ASIC1a C-terminus. TMII domains resolved by different chicken ASIC1 crystal structures are printed in blue. Unresolved flexible regions are shown in gray. Arginines in the di-arginine motif are printed in red. Note that the resolved TMII helices are of different lengths. **A2**, bottom view of the intracellular stretches and the transmembrane domain of ASIC1a channel (PDB 6AVE). The two transmembrane regions, TMI and TMII, of one subunit and the short stretches of N- and C-termini are shown in blue and yellow, respectively. **A3**, side view of the transmembrane and proximal cytoplasmic domains of superimposed closed and open states of ASIC1a channels. The resting state from PDB 6AVE is shown in gray, while the open state from PDB 4NTW is shown in orange. Note that TMII is separated in two parts, TM2a and TM2b, by a GAS belt (labeled in pink in A1). **B.** ASIC1a TMII domain sequences predicted by hydrophobicity analysis using computer algorithms listed at left. **C.** Alignment of the second transmembrane helices and proximal C-terminal domains of ASIC and ENaC family members.

**Figure S8. Surface ASIC1a and PS are both highly enriched and colocalized at the leading edge of CHO cells.**

**A.** Two more examples for Figure 5D. Representative confocal images of CHO cells transfected with N-terminal EGFP-tagged HA-ASIC1a and mCherry-Lact-C2. After fixation, cells were immunostained with anti-HA under non-permeabilized conditions to visualize surface ASIC1a (sASIC1a or s), followed by staining with anti-EGFP under permeabilized conditions to visualize total ASIC1a (tASIC1a or t) and with anti-mCherry for Lact-C2. Arrowheads indicate leading edges of the cells. White lines indicate line scan areas in (B). **B.** Intensity profiling of line scan areas through the leading edge from (A, last panel).

**Figure S9. Uncoupling PS-ASIC1a interaction alters synaptic targeting of ASIC1a.**

**A & B.** Cumulative probabilities of Pearson's correlation coefficients between total ASIC1a and Lact-C2 (A), or PSD-95 (B). n = 200 spines from 10 cells. **A.** $^{**}p$ = 0.048, one-way ANOVA with Turkey's post hoc. WT vs $^{467}$EE$^{468}$, n.s.; $^{467}$EE$^{468}$ vs $^{467}$KK$^{468}$, $^{**}p < 0.01$; WT vs $^{467}$KK$^{468}$, n.s. **B.** $^{**}p$ = 0.009, one-way ANOVA with Turkey's post hoc. WT vs $^{467}$EE$^{468}$, $^{**}p < 0.01$; $^{467}$EE$^{468}$ vs $^{467}$KK$^{468}$, n.s.; WT vs $^{467}$KK$^{468}$, n.s. **C & D.** Frequency distributions (C) and cumulative probabilities (D) of Pearson's correlation coefficients between Lact-C2 and PSD-95 for each group. n = 200 spines from 10 cells. $p$ = 0.2417, one-way ANOVA with Turkey's post hoc. WT vs $^{467}$EE$^{468}$, n.s.; $^{467}$EE$^{468}$ vs $^{467}$KK$^{468}$, n.s.; WT vs $^{467}$KK$^{468}$, n.s.

Figure S1

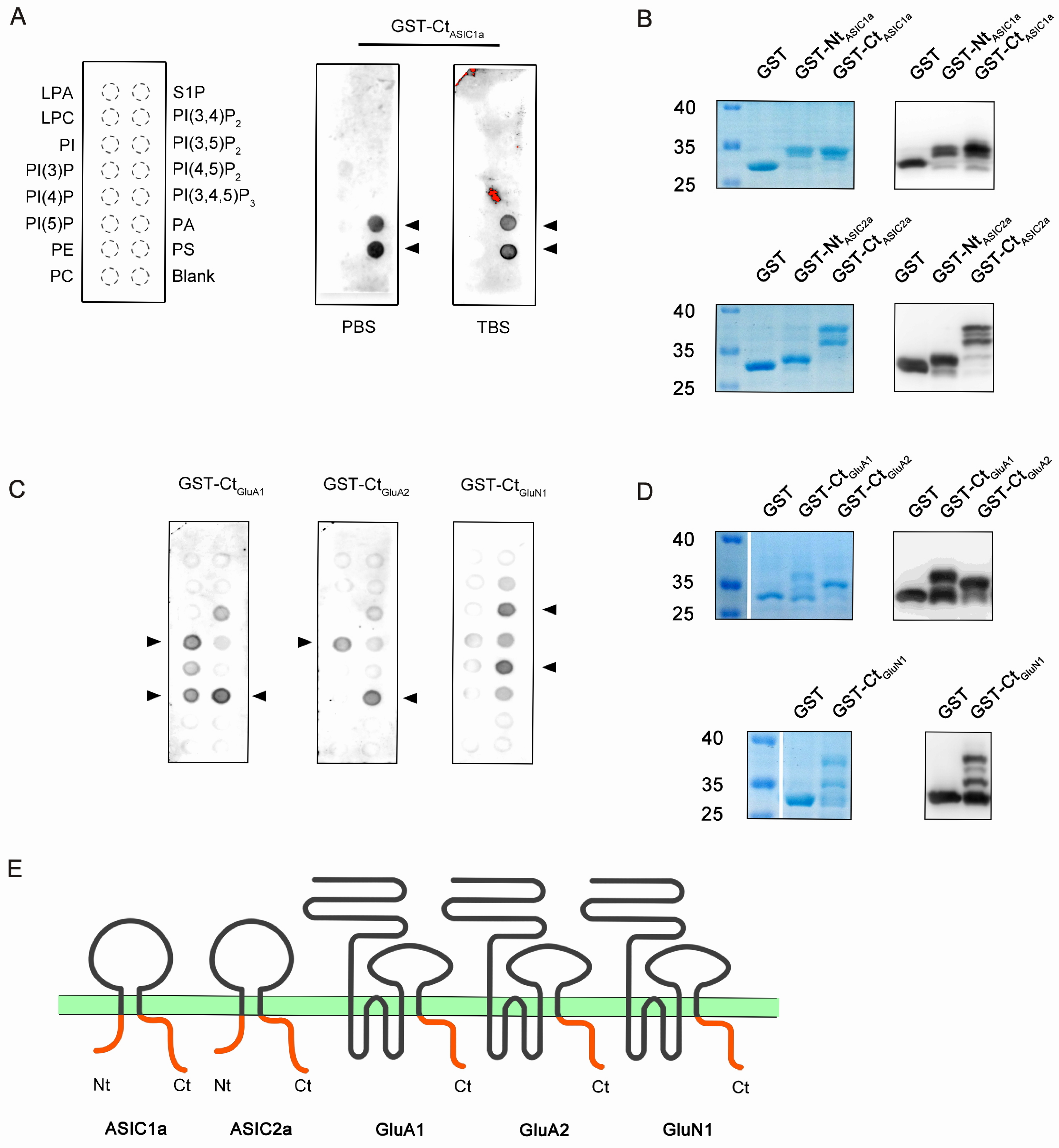

A
GST-Ct$_{ASIC1a}$
LPA
LPC
PI
PI(3)P
PI(4)P
PI(5)P
PE
PC
S1P
PI(3,4)P$_2$
PI(3,5)P$_2$
PI(4,5)P$_2$
PI(3,4,5)P$_3$
PA
PS
Blank
PBS
TBS
B
GST
GST-Nt$_{ASIC1a}$
GST-Ct$_{ASIC1a}$
40
35
25
GST-Nt$_{ASIC2a}$
GST-Ct$_{ASIC2a}$
C
GST-Ct$_{GluA1}$
GST-Ct$_{GluA2}$
GST-Ct$_{GluN1}$
D
E
Nt
Ct
ASIC1a
ASIC2a
GluA1
GluA2
GluN1

Figure S2

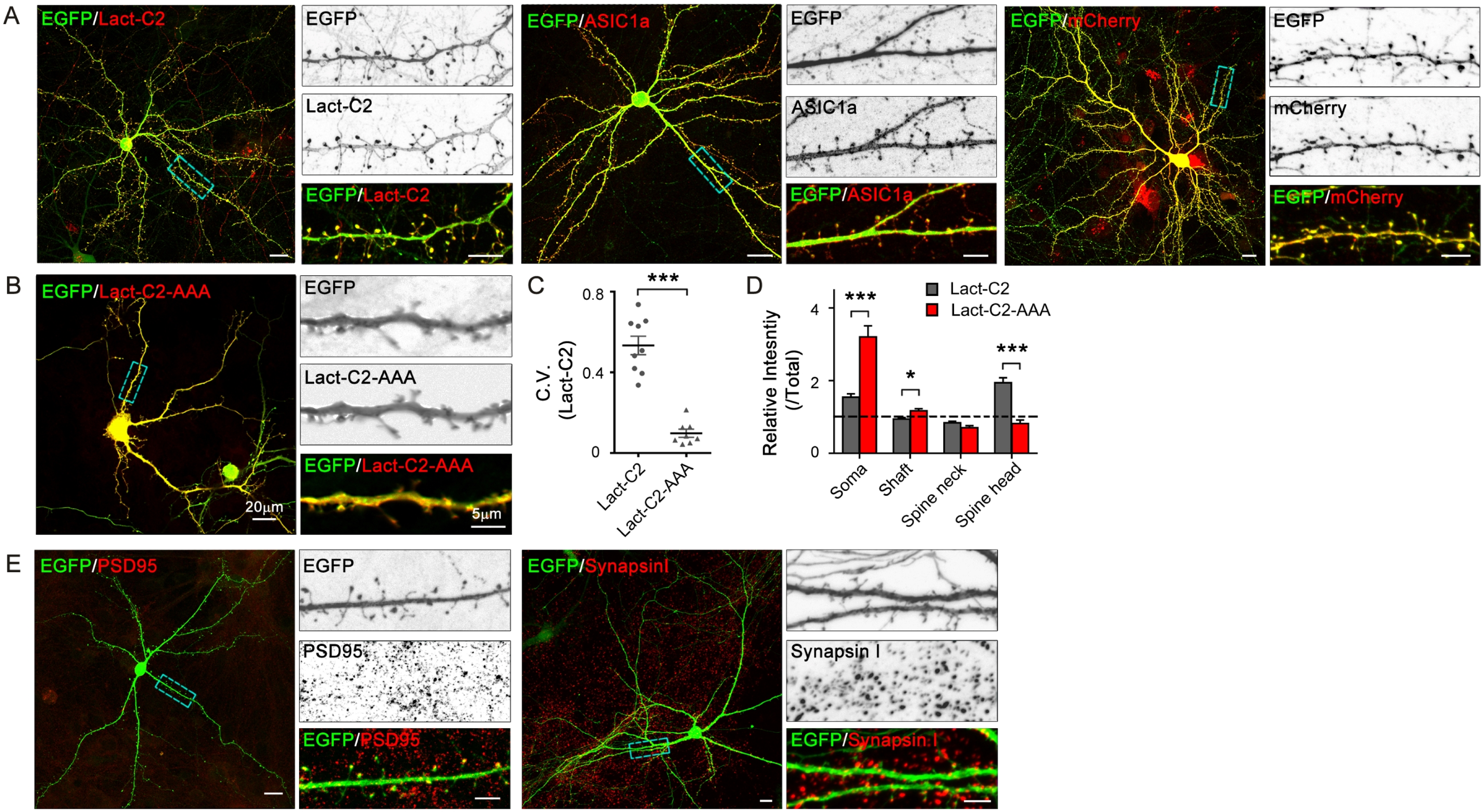
A
EGFP/Lact-C2
EGFP
Lact-C2
EGFP/Lact-C2
EGFP/ASIC1a
EGFP
ASIC1a
EGFP/ASIC1a
EGFP/mCherry
EGFP
mCherry
EGFP/mCherry
B
EGFP/Lact-C2-AAA
20μm
EGFP
Lact-C2-AAA
EGFP/Lact-C2-AAA
5μm
C
C.V. (Lact-C2)
0.8
0.4
0
***
Lact-C2
Lact-C2-AAA
D
Relative Intesntiy (/Total)
4
2
0
Lact-C2
Lact-C2-AAA
***
*
***
Soma
Shaft
Spine neck
Spine head
E
EGFP/PSD95
EGFP
PSD95
EGFP/PSD95
EGFP/SynapsinI
Synapsin I
EGFP/Synapsin I

Figure S3

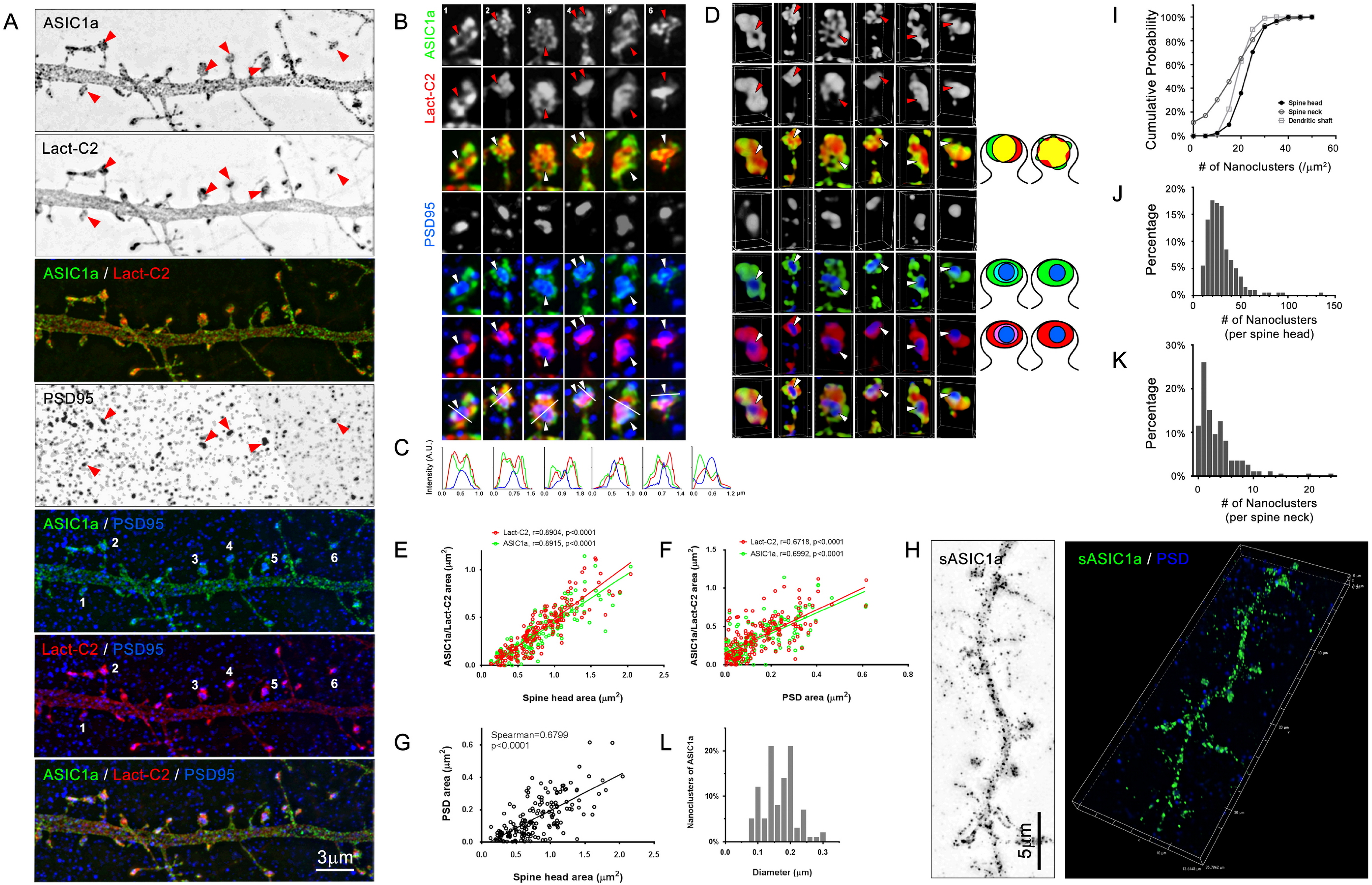
A
ASIC1a
Lact-C2
ASIC1a / Lact-C2
PSD95
ASIC1a / PSD95
Lact-C2 / PSD95
ASIC1a / Lact-C2 / PSD95
3μm
B
ASIC1a
Lact-C2
PSD95
C
Intensity (A.U.)
D
E
Lact-C2, r=0.8904, p<0.0001
ASIC1a, r=0.8915, p<0.0001
ASIC1a/Lact-C2 area (μm²)
Spine head area (μm²)
F
Lact-C2, r=0.6718, p<0.0001
ASIC1a, r=0.6992, p<0.0001
PSD area (μm²)
G
Spearman=0.6799
p<0.0001
PSD area (μm²)
Spine head area (μm²)
L
Nanoclusters of ASIC1a
Diameter (μm)
H
sASIC1a
5μm
sASIC1a / PSD
I
Cumulative Probability
Spine head
Spine neck
Dendritic shaft
### of Nanoclusters (/μm²)
J
Percentage
### of Nanoclusters
(per spine head)
K
Percentage
### of Nanoclusters
(per spine neck)

Figure S4

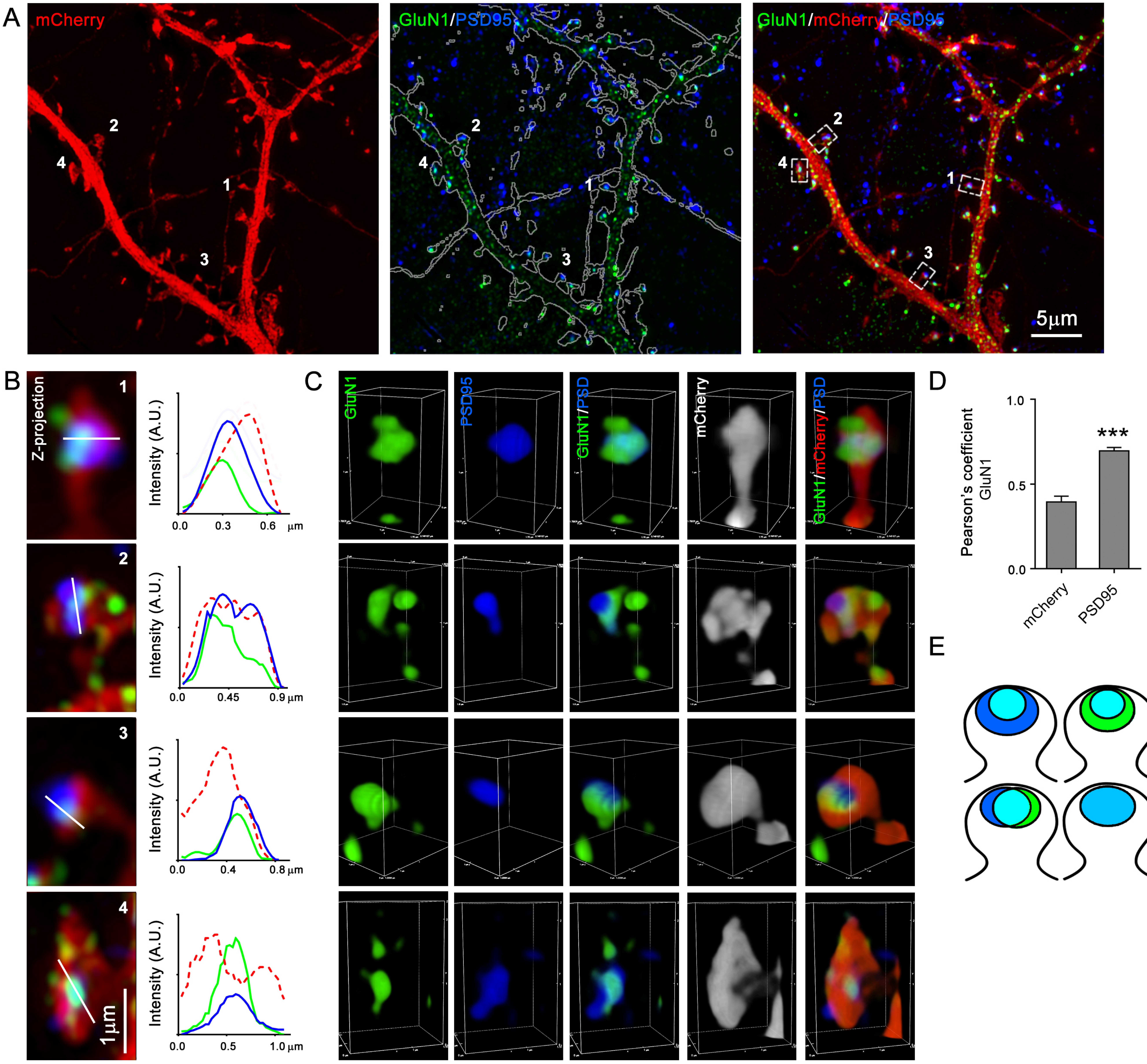
A
mCherry
GluN1/PSD95
GluN1/mCherry/PSD95
1
2
3
4
5μm
B
Z-projection
Intensity (A.U.)
0.0
0.3
0.6
μm
0.45
0.9 μm
0.4
0.8 μm
0.5
1.0 μm
1μm
C
GluN1
PSD95
GluN1/PSD
mCherry
GluN1/mCherry/PSD
D
Pearson's coefficient
GluN1
1.0
0.5
0.0
***
mCherry
PSD95
E

#### Figure S5

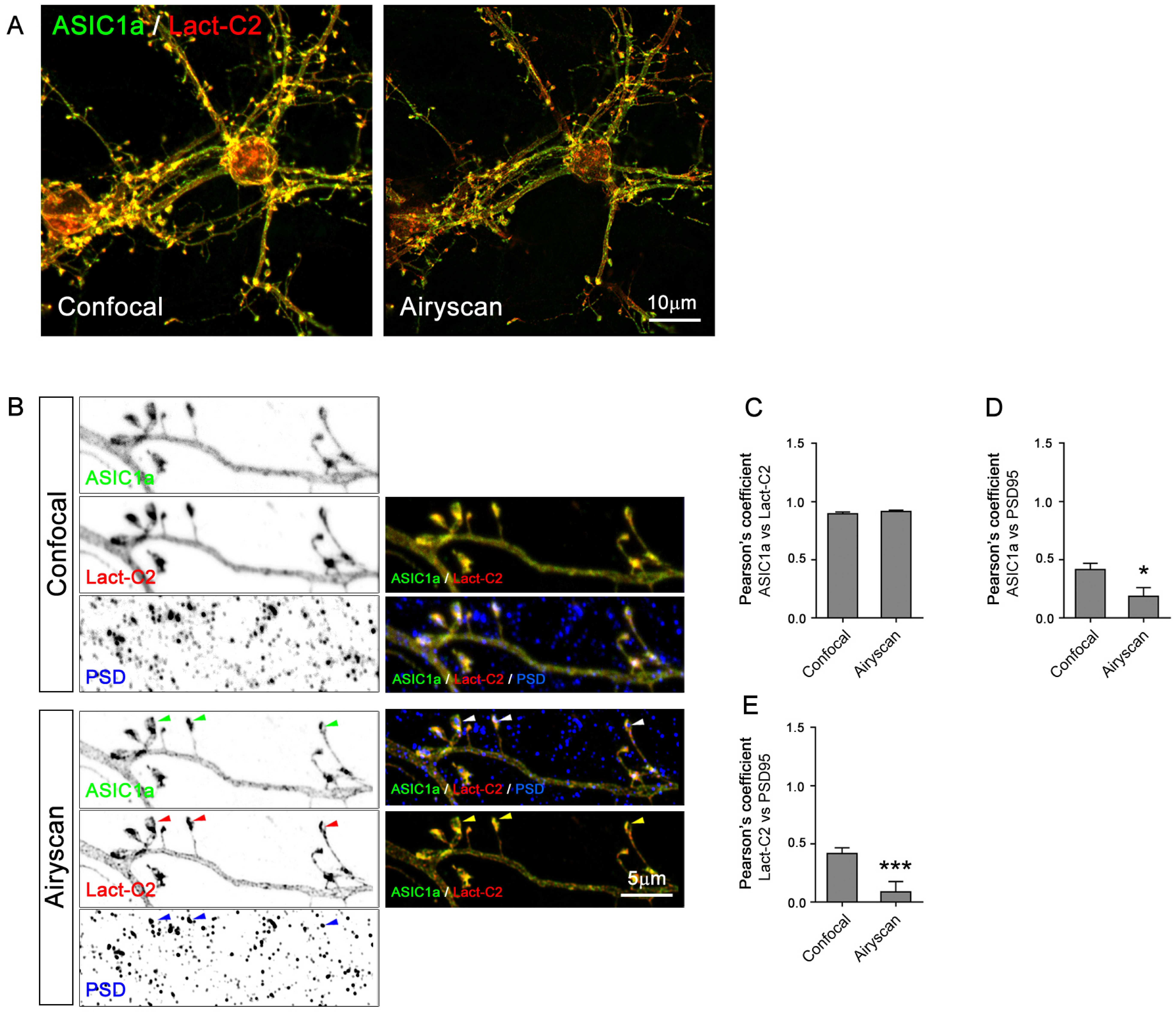
A
ASIC1a / Lact-C2
Confocal
Airyscan
10µm
B
Confocal
ASIC1a
Lact-C2
PSD
ASIC1a / Lact-C2
ASIC1a / Lact-C2 / PSD
Airyscan
ASIC1a
Lact-C2
PSD
ASIC1a / Lact-C2 / PSD
ASIC1a / Lact-C2
5µm
C
Pearson's coefficient
ASIC1a vs Lact-C2
1.5
1.0
0.5
0.0
Confocal
Airyscan
D
Pearson's coefficient
ASIC1a vs PSD95
1.5
1.0
0.5
0.0
*
Confocal
Airyscan
E
Pearson's coefficient
Lact-C2 vs PSD95
1.5
1.0
0.5
0.0
***
Confocal
Airyscan

Figure S6

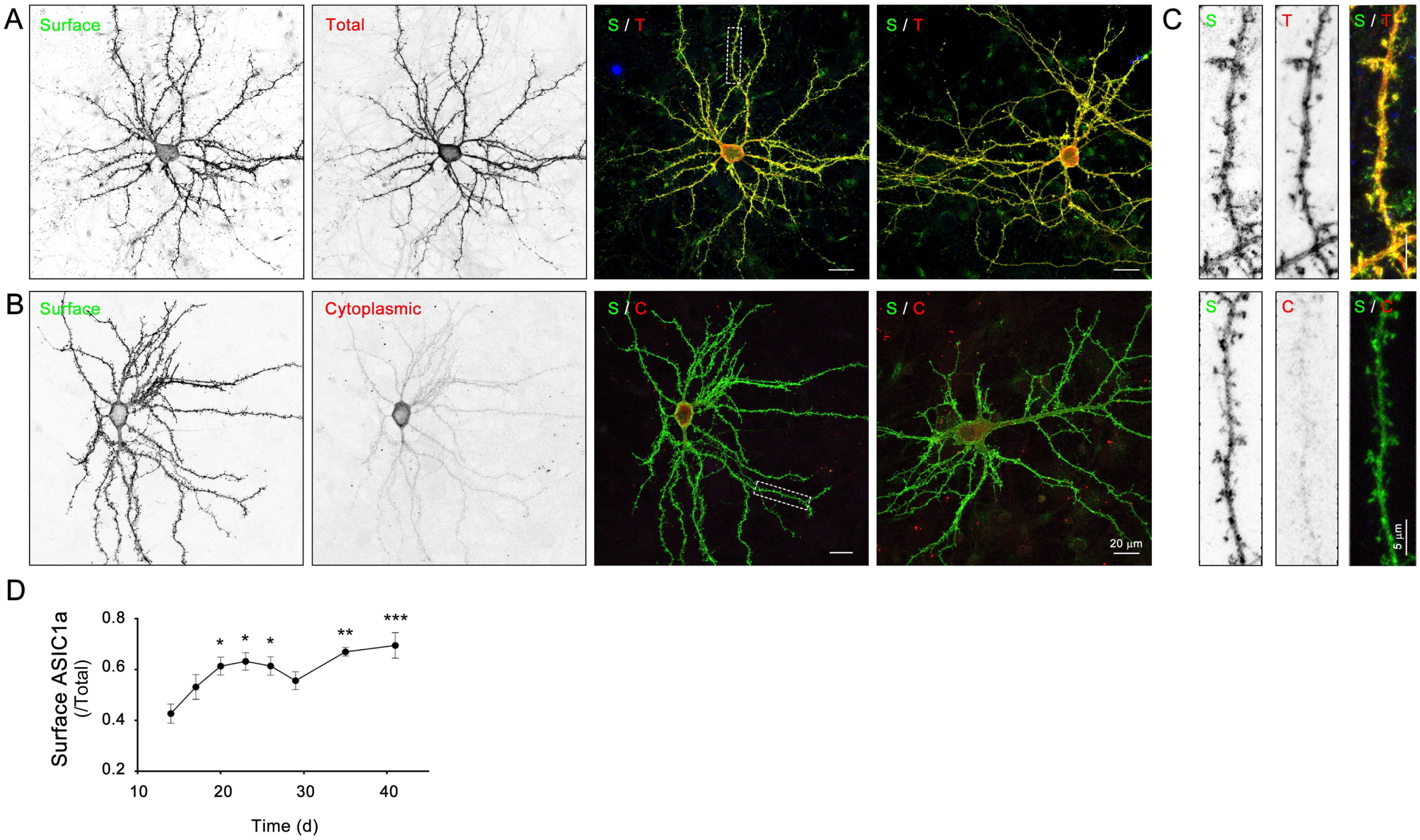

A
Surface
Total
S / T
S / T
C
S
T
S / T
B
Surface
Cytoplasmic
S / C
S / C
20 µm
S
C
S / C
5 µm
D
Surface ASIC1a (/Total)
0.8
0.6
0.4
0.2
10
20
30
40
Time (d)
*
*
*
**
***

Figure S7

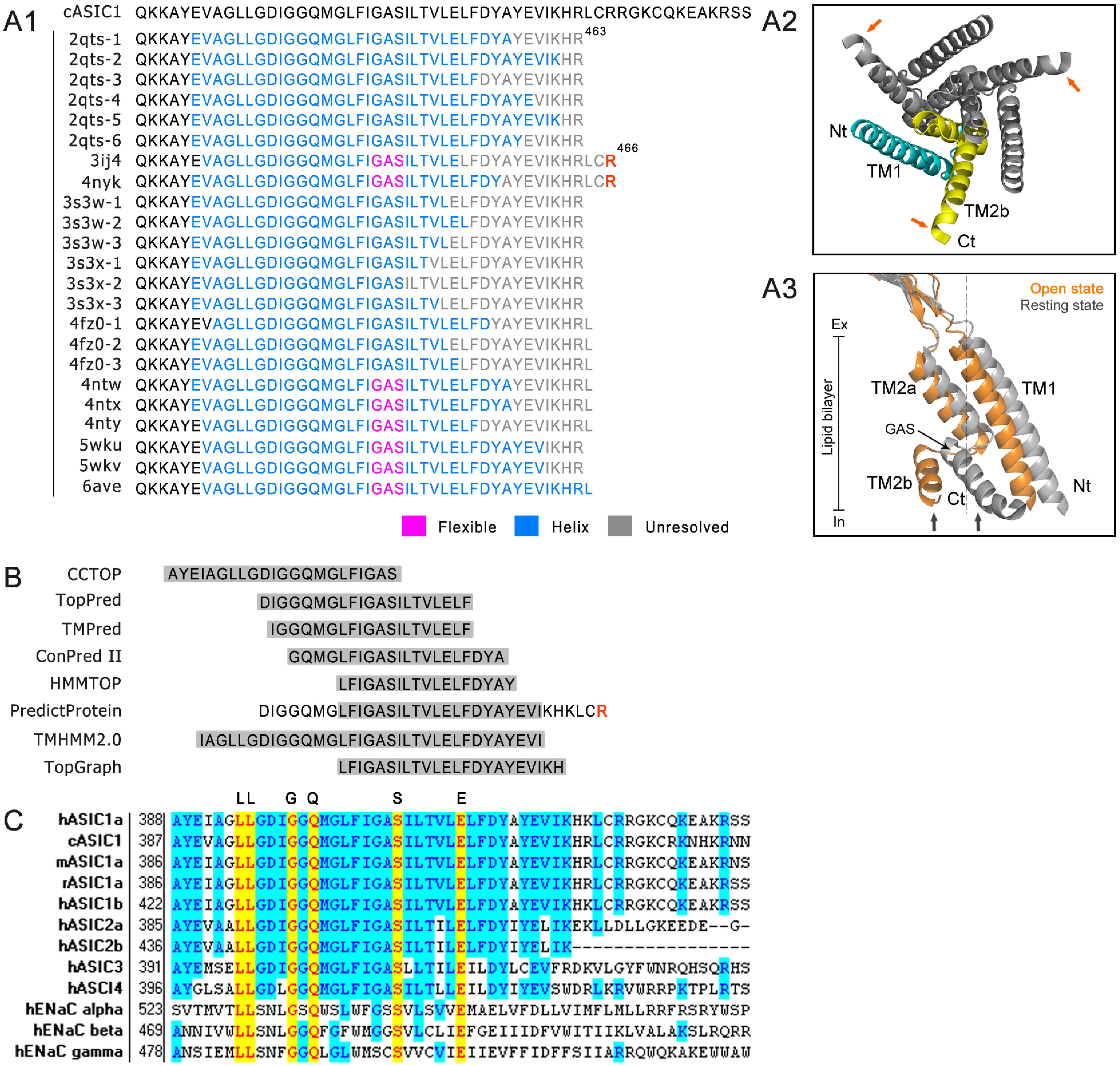

A1
cASIC1 QKKAYEVAGLLGDIGGQMGLFIGASILTVLELFDYAYEVIKHRLCRRGKCQKEAKRSS
2qts-1 QKKAYEVAGLLGDIGGQMGLFIGASILTVLELFDYAYEVIKHR 463
2qts-2 QKKAYEVAGLLGDIGGQMGLFIGASILTVLELFDYAYEVIKHR
2qts-3 QKKAYEVAGLLGDIGGQMGLFIGASILTVLELFDYAYEVIKHR
2qts-4 QKKAYEVAGLLGDIGGQMGLFIGASILTVLELFDYAYEVIKHR
2qts-5 QKKAYEVAGLLGDIGGQMGLFIGASILTVLELFDYAYEVIKHR
2qts-6 QKKAYEVAGLLGDIGGQMGLFIGASILTVLELFDYAYEVIKHR
3ij4 QKKAYEVAGLLGDIGGQMGLFIGASILTVLELFDYAYEVIKHRLCR 466
4nyk QKKAYEVAGLLGDIGGQMGLFIGASILTVLELFDYAYEVIKHRLCR
3s3w-1 QKKAYEVAGLLGDIGGQMGLFIGASILTVLELFDYAYEVIKHR
3s3w-2 QKKAYEVAGLLGDIGGQMGLFIGASILTVLELFDYAYEVIKHR
3s3w-3 QKKAYEVAGLLGDIGGQMGLFIGASILTVLELFDYAYEVIKHR
3s3x-1 QKKAYEVAGLLGDIGGQMGLFIGASILTVLELFDYAYEVIKHR
3s3x-2 QKKAYEVAGLLGDIGGQMGLFIGASILTVLELFDYAYEVIKHR
3s3x-3 QKKAYEVAGLLGDIGGQMGLFIGASILTVLELFDYAYEVIKHR
4fz0-1 QKKAYEVAGLLGDIGGQMGLFIGASILTVLELFDYAYEVIKHRL
4fz0-2 QKKAYEVAGLLGDIGGQMGLFIGASILTVLELFDYAYEVIKHRL
4fz0-3 QKKAYEVAGLLGDIGGQMGLFIGASILTVLELFDYAYEVIKHRL
4ntw QKKAYEVAGLLGDIGGQMGLFIGASILTVLELFDYAYEVIKHRL
4ntx QKKAYEVAGLLGDIGGQMGLFIGASILTVLELFDYAYEVIKHRL
4nty QKKAYEVAGLLGDIGGQMGLFIGASILTVLELFDYAYEVIKHRL
5wku QKKAYEVAGLLGDIGGQMGLFIGASILTVLELFDYAYEVIKHR
5wkv QKKAYEVAGLLGDIGGQMGLFIGASILTVLELFDYAYEVIKHR
6ave QKKAYEVAGLLGDIGGQMGLFIGASILTVLELFDYAYEVIKHRL
Flexible
Helix
Unresolved
A2
Nt
TM1
TM2b
Ct
A3
Open state
Resting state
Ex
Lipid bilayer
In
TM2a
TM1
GAS
TM2b
Ct
Nt
B
CCTOP AYEIAGLLGDIGGQMGLFIGAS
TopPred DIGGQMGLFIGASILTVLELF
TMPred IGGQMGLFIGASILTVLELF
ConPred II GQMGLFIGASILTVLELFDYA
HMMTOP LFIGASILTVLELFDYAY
PredictProtein DIGGQMGLFIGASILTVLELFDYAYEVIKHKLCR
TMHMM2.0 IAGLLGDIGGQMGLFIGASILTVLELFDYAYEVI
TopGraph LFIGASILTVLELFDYAYEVIKH
C
LL G Q S E
hASIC1a 388 AYEIAGLLGDIGGQMGLFIGASILTVLELFDYAYEVIKHKLCRRGKCQKEAKRSS
cASIC1 387 AYEVAGLLGDIGGQMGLFIGASILTVLELFDYAYEVIKHRLCRRGKCRKNHKRNN
mASIC1a 386 AYEIAGLLGDIGGQMGLFIGASILTVLELFDYAYEVIKHRLCRRGKCQKEAKRNS
rASIC1a 386 AYEIAGLLGDIGGQMGLFIGASILTVLELFDYAYEVIKHRLCRRGKCQKEAKRSS
hASIC1b 422 AYEIAGLLGDIGGQMGLFIGASILTVLELFDYAYEVIKHKLCRRGKCQKEAKRSS
hASIC2a 385 AYEVAALLGDIGGQMGLFIGASILTILELFDYIYELIKEKLLDLLGKEEDE--G-
hASIC2b 436 AYEVAALLGDIGGQMGLFIGASILTILELFDYIYELIK-----------------
hASIC3 391 AYEMSELLGDIGGQMGLFIGASLLTILEILDYLCEVFRDKVLGYFWNRQHSQRHS
hASCI4 396 AYGLSALLGDLGGQMGLFIGASILTLLEILDYIYEVSWDRLKRVWRRPKTPLRTS
hENaC alpha 523 SVTMVTLLSNLGSQWSLWFGSSVLSVVEMAELVFDLLVIMFLMLLRRFRSRYWSP
hENaC beta 469 ANNIVWLLSNLGGQFGFWMGGSVLCLIEFGEIIIDFVWITIIKLVALAKSLRQRR
hENaC gamma 478 ANSIEMLLSNFGGQLGLWMSCSVVCVIEIIEVFFIDFFSIIARRQWQKAKEWWAW

**Figure S8**

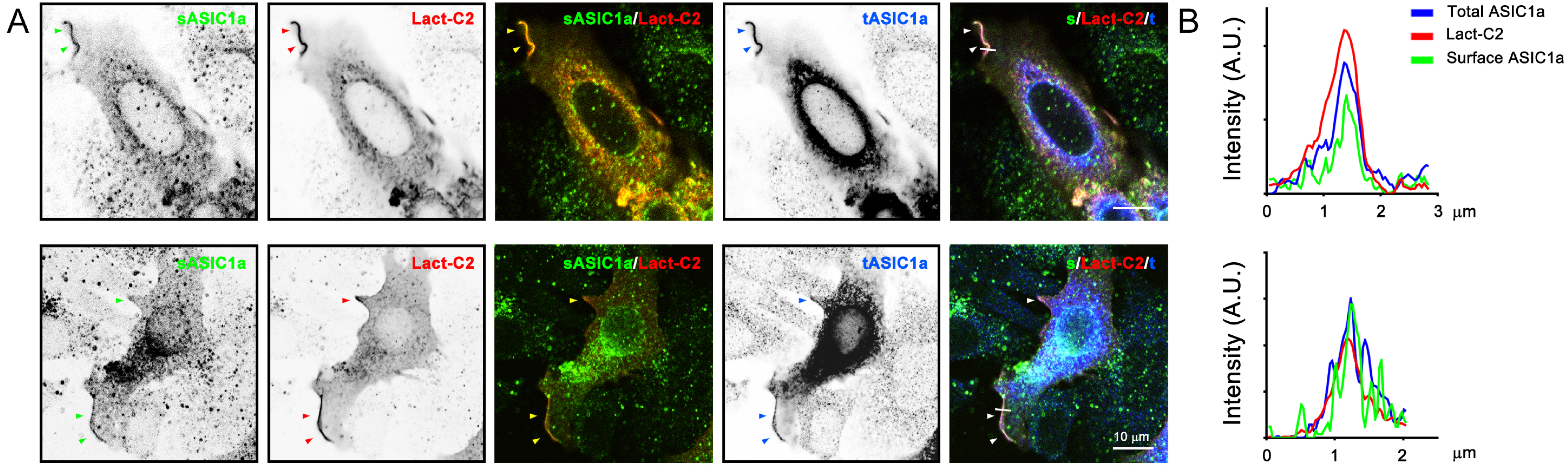
A
sASIC1a
Lact-C2
sASIC1a/Lact-C2
tASIC1a
s/Lact-C2/t
10 µm
B
Total ASIC1a
Lact-C2
Surface ASIC1a
Intensity (A.U.)
0
1
2
3
µm

Figure S9

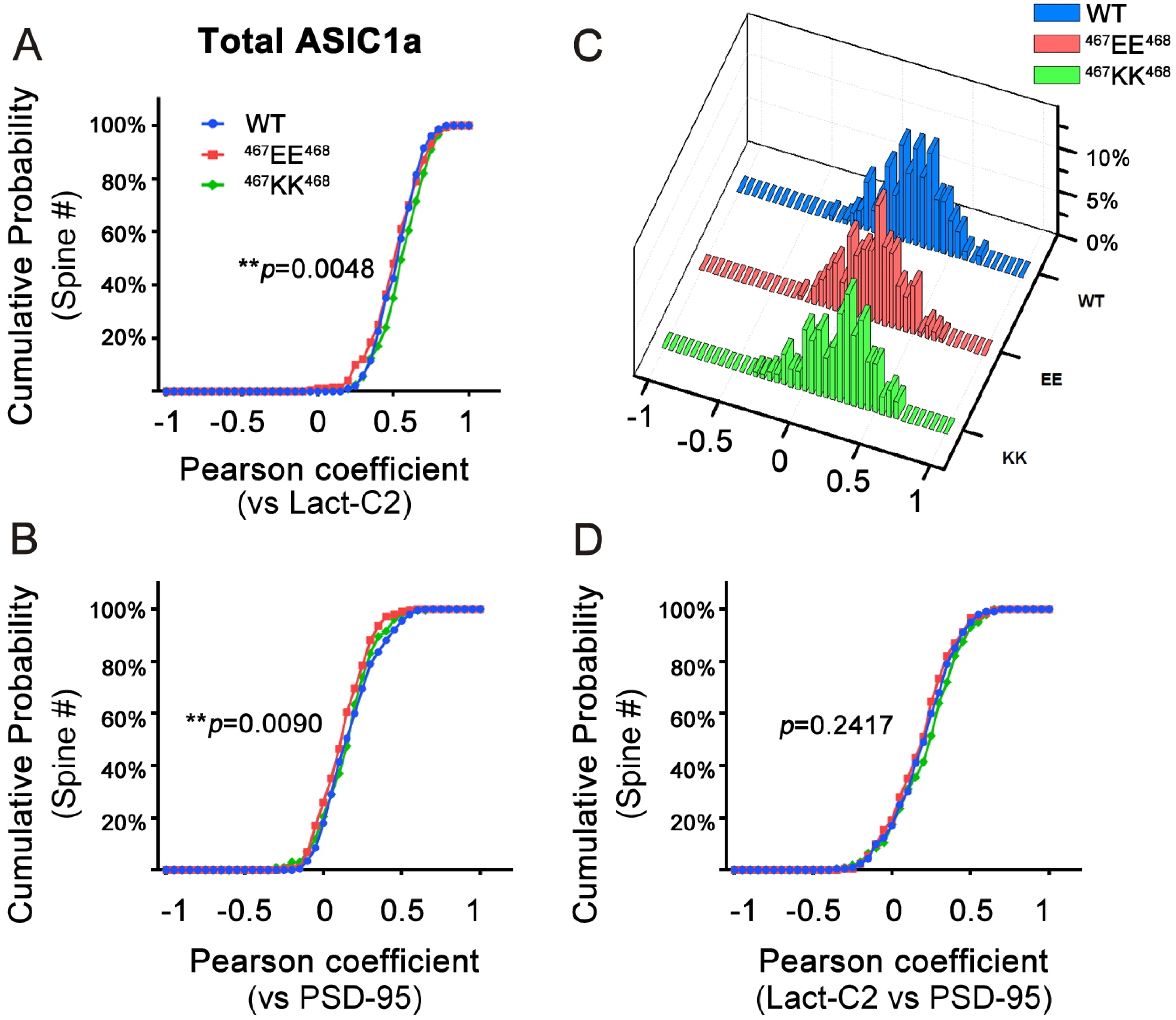
A
Total ASIC1a
Cumulative Probability
(Spine #)
100%
80%
60%
40%
20%
WT
467EE468
467KK468
**p=0.0048
-1
-0.5
0
0.5
1
Pearson coefficient
(vs Lact-C2)
C
WT
467EE468
467KK468
10%
5%
0%
WT
EE
KK
-1
-0.5
0
0.5
1
B
Cumulative Probability
(Spine #)
100%
80%
60%
40%
20%
**p=0.0090
-1
-0.5
0
0.5
1
Pearson coefficient
(vs PSD-95)
D
Cumulative Probability
(Spine #)
100%
80%
60%
40%
20%
p=0.2417
-1
-0.5
0
0.5
1
Pearson coefficient
(Lact-C2 vs PSD-95)
